# Beyond reference bias: Making pangenomes accessible with PangyPlot

**DOI:** 10.1101/2025.10.31.684064

**Authors:** Scott Mastromatteo, Shalvi Chirmade, Delnaz Roshandel, Bhooma Thiruvahindrapuram, Zhuozhi Wang, Rohan V Patel, Wilson WL Sung, Amirhossein Hajianpour, Cheng Wang, Fan Lin, Katherine Keenan, Julie Avolio, Paul Eckford, Felix Ratjen, Canadian Cystic Fibrosis Gene Modifier Consortium, Lisa J Strug

## Abstract

Linear reference genomes have standardized genomics research but remain limited by reference bias, which skews read mapping and variant discovery. This bias can distort the interpretation of genetic variation, particularly for populations that are genetically distant from the reference. Pangenome graphs, such as those generated by the Human Pangenome Reference Consortium (HPRC), mitigate this limitation by integrating diverse haplotypes into a unified representation of human genetic variation. However, the complexity of graph-based data and the lack of intuitive visualization tools have hindered broader adoption.

Here we introduce *PangyPlot*, a genome browser that simplifies exploration of pangenome graphs by retaining linear-style navigation, integrating gene annotations, abstracting complex variation into interpretable structures, and employing a dynamic, physics-based layout optimization engine. We demonstrate its utility by constructing a chromosome 7 graph from 101 individuals with cystic fibrosis (CF), capturing a broad spectrum of genetic variation. Using *PangyPlot*, we visualized CF-relevant loci and compared results with existing graph viewers, highlighting its ability to display both base-level and large structural variation. With an additional 64 PacBio HiFi assemblies, we fine-mapped a repeat-dense CF modifier locus on chromosome 5, where *PangyPlot* was used in conjunction with graph-based analysis to identify a repeat expansion in the 5^***′***^ end of *EXOC3* that may promote G-quadruplex formation and affect gene expression. Together, these examples demonstrate *PangyPlot* ‘s capacity to make population-level variation interpretable. To support broader use of graph-based resources, we also released a live public instance of *PangyPlot* preloaded with HPRC data (https://pangyplot.research.sickkids.ca/).

## 1 Introduction

Genome sequencing studies typically transform raw sequence reads into alignments against an established reference genome to facilitate research of gene regulation, function, and disease association. The reference genome provides a unified coordinate system and enables consistent comparison of an individual’s genome to public databases of genetic variation and annotations. Such comparisons rely on accurate mapping of reads to the reference, which can sometimes be unreliable. Ambiguous read mapping occurs when the read contains insufficient information to map uniquely to the reference, a well-known limitation with short-read sequencing [1]. Low mappability is particularly problematic in regions characterized by long stretches of low sequence complexity or repetitive DNA such as copy number variation (CNV) and variable number of tandem repeats (VNTR). Long-read sequencing technologies provided by Oxford Nanopore Technologies (ONT) and Pacific Biosciences (PacBio) partially mitigate these issues by generating longer reads with significantly improved mappability [2].

In contrast to advances in sequencing technology, the reference genome model established during the Human Genome Project in 2001 [3] has remained largely unchanged. While subsequent builds from the Genome Reference Consortium (GRCh37, GRCh38) and the Telomere-to-Telomere (T2T) assembly of CHM13 [4] represent notable improvements in the completeness and accuracy of the human reference genome, they are all fundamentally linear references: contiguous string representations of a single haploid sequence. Linear reference genomes are susceptible to reference bias, where differences between the reads and the reference decrease mappability and lead to systematic undercalling of alternative variation [5]. Reference bias disproportionately affects populations whose genomes differ significantly from the reference, potentially exacerbating disparities in genetic research and clinical interpretation. GRCh38 attempts to mitigate reference bias by including a small number of alternative contigs for highly variable loci [6], but these contigs are often omitted from analyses because they complicate alignment and interpretation.

To better represent the full range of diversity in the human genome, pangenomes have been proposed as an alternative to standard linear references [7]. Pangenomes use information from multiple individuals to create a unified representation of all observed variation—from small single-nucleotide variants to large structural variants. Better representation of complex loci reduces reference bias and improves read mapping and variant calling. The Human Pangenome Reference Consortium (HPRC) has generated human reference pangenomes [8] constructed by combining ancestrally diverse *de novo* assemblies into variation graphs. In these graphs, DNA sequences are represented by nodes, with variation encoded in edges (Fig. 1a–c). Variation graphs commonly form bubbles (also known as snarls), in which a set of connected nodes shares a single source and sink and can form complex, nested structures [9] (Fig. 1d,e). These higher-order structures can be identified using existing algorithms [10]. Since variants are encoded directly in graph topology, understanding pangenome structure can help improve the interpretation and discovery of complex genetic variation.

**Fig. 1.**
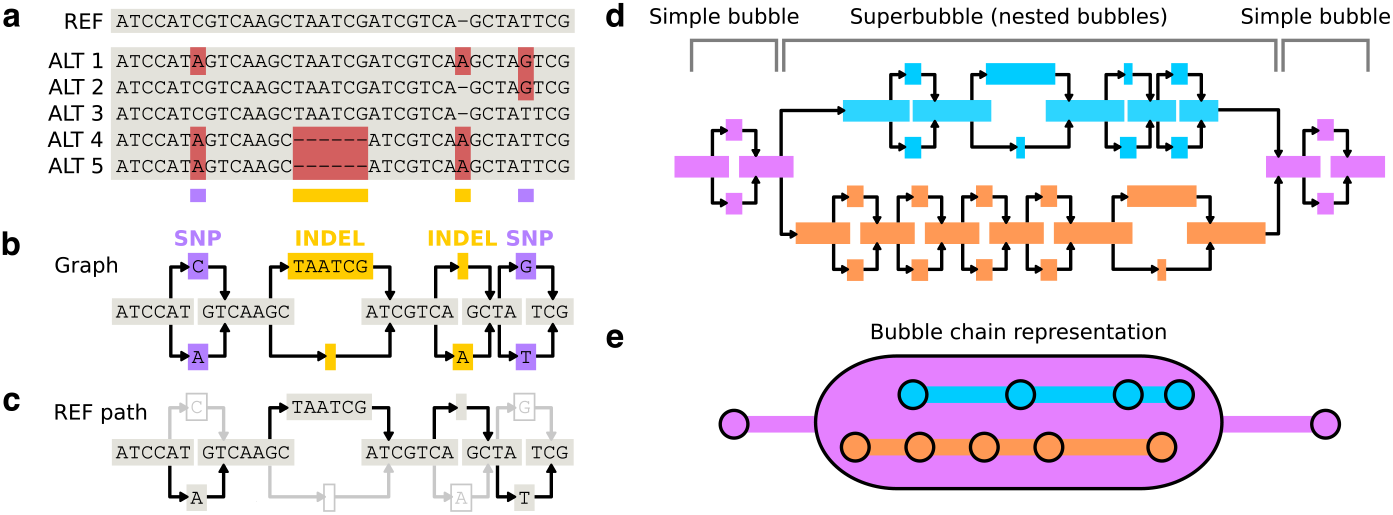
Modeling genetic variation using graph-based structures. **a**, Linear reference sequence with example haplotypes aligned. Differences relative to the reference are highlighted in red. **b**, Graph-based representation of the haplotypes. Each of the four biallelic variants is represented as a bubble: a source node with bifurcating edges to two allele nodes and converging on a single sink node. **c**, A specific haplotype can be defined as a path through the graph. Reconstruction of the reference sequence is shown. **d**, Example of a complex graph structure. Purple nodes form simple biallelic bubbles, while orange and blue nodes represent two branches of a superbubble containing nested bubbles. **e**, Bubble chains formed from adjacent bubbles. Three bubble chains are shown; circles denote bubbles linked in a chain. The purple chain consists of two simple bubbles and a superbubble, while the nested blue and orange branches define separate bubble chains within the superbubble.

The graph construction method used is important to topology [11]. The HPRC v1.1 pangenome release includes graphs created by three different tools. Minigraph [12] uses an iterative approach of starting from a single assembly and iteratively aligning one assembly at a time while incorporating structural variants as bubbles. The resulting graph, however, is order-dependent and biased toward the first assembly [13]. Minigraph-Cactus [14] is an extension that adds base-level alignment and graph pruning. Pruning methods include `full` (no pruning), `clip` (removal of large unaligned or dangling nodes), and `filter` (only includes nodes supported by a minimum number of haplotypes). The PanGenome Graph Builder (`PGGB`) [15] relies on an all-to-all alignment approach that is unbiased but scales poorly as the number of assemblies grows.

Once built, the graph can be stored in many different file formats. The commonly used Graphical Fragment Assembly (GFA) format represents graphs as a series of segment (`S`) and link (`L`) lines, with path (`P`) or walk (`W`) lines depending on the specification (GFA 1.0, GFA 2.0, GFA 1.1). However, the plain-text GFA format is insufficient for many graph operations. Specialized toolkits rely on more optimized file formats: the variation graph toolkit (`vg`) [16] encodes variation-aware structures in indexed binary files, and `odgi` [17] uses a format optimized for large-scale graph manipulation. The same graph is often converted between formats to access specific operations, which requires familiarity with multiple file formats and toolkits. Furthermore, in contrast to linear references that provide a simple coordinate system, graph coordinates are path-dependent and unstable, as incorporation of new variation can disrupt or shift existing coordinates.

Human-scale pangenome visualization also remains difficult, owing to the massive scale of the data (hundreds of millions of nodes and edges) and the complexity of producing an interpretable visual layout. The most widely used sequence graph visualization tool, Bandage [18], was originally developed for *de novo* assembly graphs in GFA format. While it has been adapted for variation graphs, their substantially greater complexity exposes key limitations: Bandage loads the entire graph into memory and computes layouts dynamically, restricting the size and complexity of graphs that can be effectively visualized. Another visualization tool, Sequence Tube Map [19], renders small portions of `vg` variation graphs by drawing paths as interweaving lines. In contrast, the `odgi` toolkit includes a GPU-accelerated two-dimensional layout algorithm [20] and is capable of rendering static images of complete graphs (`odgi viz` for one dimension and `odgi draw` for two dimensions), but it does not support interactive exploration or dynamic visualization. As a result, what should be the simple task of visualizing a region of interest in public pangenome data becomes prohibitively complex. The workflow requires users to choose between different graph construction approaches, extract subgraphs by coordinates, and often perform multiple commandline file conversions before loading the visualization software. Many other tools have attempted to address the visualization problem [21–25], yet a gap remains in the general accessibility and usability of pangenomic resources such as the HPRC data.

To address this gap, we developed *PangyPlot*, a web-based reference pangenome browser designed for interactive exploration of variation graphs using familiar coordinate-based navigation inspired by linear reference genomes. *PangyPlot* can either be hosted on a server as a public utility or run locally with custom pangenome data for research and clinical applications. Here, we demonstrate both by making a public HPRC v1.1 resource available at https://pangyplot.research.sickkids.ca/ and by visualizing variation graphs we constructed from individuals participating in the Canadian Cystic Fibrosis (CF) Gene Modifier Study. Using all of our available data, we generated 101 assemblies with a hybrid approach combining PacBio continuous long read (CLR) and 10x Genomics (10XG) linked reads, as well as 64 assemblies from PacBio HiFi data. Given that CF is more common in individuals of European ancestry [26], we assessed how an ancestrally homogeneous cohort impacts genetic diversity representation in a variation graph, in contrast to the ancestrally diverse HPRC cohort. We then used *PangyPlot* to visualize three loci relevant to CF: (*i*) *CFTR*, which harbors over 1,000 CF-causing variants including complex alleles [27]; (*ii*) the *PRSS1–PRSS2* trypsinogen locus, which contains a CNV previously associated with intestinal obstruction at birth in CF (meconium ileus, MI) [28, 29] and with pancreatitis risk in non-CF individuals [30]; and (*iii*) a chromosome 5 subtelomeric locus (chr5p15.33) associated with CF lung disease [31, 32] and enriched for repetitive elements.

## 2 Results

### 2.1 A Canadian CF variation graph captures a broad range of genetic diversity

To enable high-quality graph construction, we first generated hybrid *de novo* assemblies (integrating CLR and 10XG reads from the same individual; detailed in Supplementary Methods 1) to produce phased assemblies for 101 individuals with CF and public sample HG002. These assemblies were then used as input to Minigraph-Cactus [14] to construct three chromosome 7 variation graphs under different pruning settings. Table 1 summarizes these graphs alongside the corresponding HPRC chromosome 7 graphs. The HPRC `full` graph contained 424 Mb of sequence—nearly double the size of the CF `full` graph and 2.6× longer than the CHM13 chr7 contig (160.6 Mb)—suggesting potential redundancy in the HPRC graph. In contrast, the CF and HPRC `clip` graphs showed similar and reasonable cumulative sequence lengths. Notably, the `d9-filter` graphs both had approximately 2.4 Mb more sequence relative to CHM13 chr7 (and 3.1 Mb more than GRCh38).

**Table 1.**
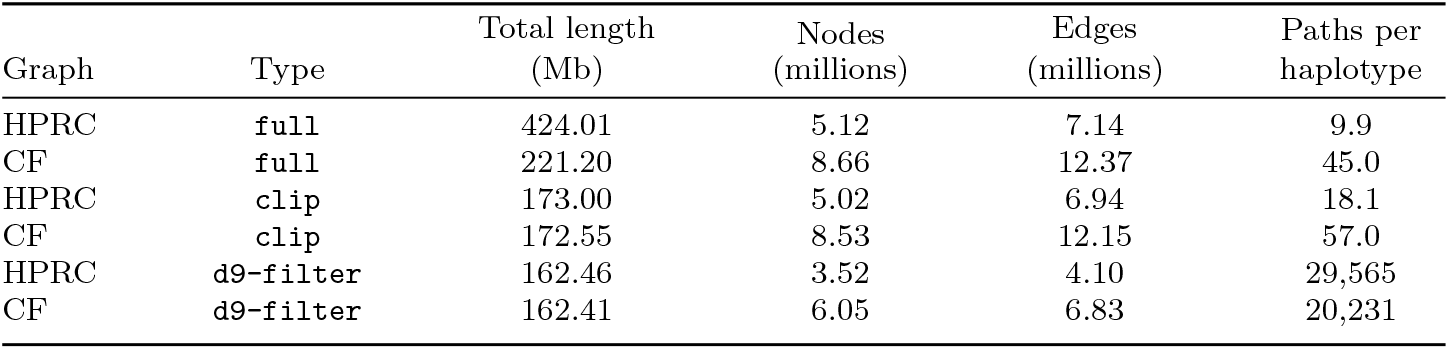
Chromosome 7 sequence graph statistics. Comparison of chromosome 7 graphs from the HPRC v1.1 release (44 samples plus GRCh38 and CHM13) and those constructed from the CF cohort (101 samples plus HG002, GRCh38, CHM13, and nine GRCh38 alternative contigs). Three graphs were generated: `full` (no pruning), `clip` (removal of large artifacts), and `d9-filter` (nodes supported by at least nine haplotypes). Metrics include cumulative sequence length, node and edge counts, and average paths per haplotype (HPRC: 90 complete haplotypes; `CF: 206` complete haplotypes). In the `full` graph, each path corresponds one-to-one with an assembled scaffold, whereas the clip and `d9-filter` pruning steps split a path whenever nodes along it are removed.

We assessed the ancestral composition and variant diversity captured by our CFspecific variation graph. Genome-wide ancestry analysis of the 101 samples revealed a predominance of individuals with European ancestry (*n* = 94), consistent with expectations for a North American CF cohort [33]. We also identified individuals with Latin American, South Asian, African American, and African ancestries (Fig. 2a). This distribution is representative of our larger short-read Canadian CF cohort (*n* = 564), in which 11.3% of individuals had non-European ancestry (Supplementary Fig. 8). In contrast, the HPRC cohort (*n* = 44) was designed to emphasize global genetic diversity (51% African, 34% Americas, 13% Asian [8]; Fig. 2b).

**Fig. 2.**
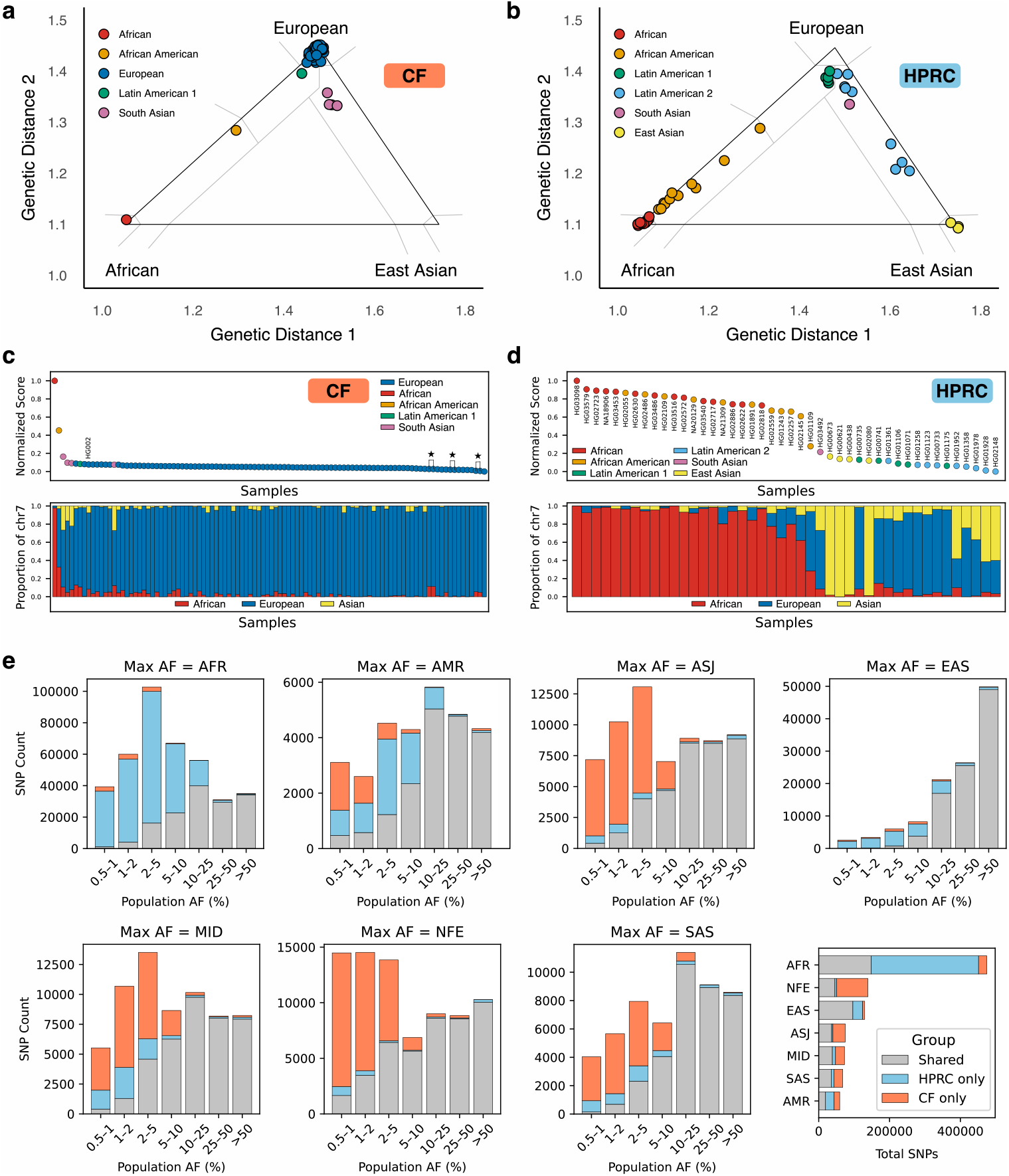
HPRC and CF cohorts display distinct ancestral compositions. Ancestry was estimated using GRAF-pop [35] for the **a**, CF cohort (94 European, 1 African, 1 African American, 1 Latin American I, 4 South Asian) and **b**, HPRC cohort (14 African, 11 African American, 4 East Asian, 5 Latin American I, 9 Latin American II, 1 South Asian). Normalized DI scores—reflecting each individual’s relative contribution to the genetic diversity of the graph—are shown for the **c**, CF and **d**, HPRC graphs. Points are colored by ancestry; publicly available samples are labeled, and sibling pairs are indicated by stars. Lower panels show detailed chromosome 7 ancestry breakdowns for each sample (African in red, European in blue, Asian in yellow), ordered from highest to lowest scores. **e**, Chromosome 7 SNVs called from CF and HPRC graphs annotated using gnomAD allele frequencies. SNVs were grouped by population with the highest allele frequency (*popmax* categories: AFR, AMR, ASJ, EAS, MID, NFE, SAS) and subdivided by allele frequency bins. Each bin is colored by the proportion of SNVs found exclusively in the CF graph (orange), exclusively in the HPRC graph (blue), or in both graphs (grey).

We devised a quantitative metric, the Diversity Importance (DI) score (see Methods), to evaluate how much each individual contributes to genetic diversity in the variation graph. Individuals carrying unique or uncommon variation relative to others in the graph receive higher scores. In the CF cohort, individuals with non-European ancestral backgrounds generally had higher DI scores (Fig. 2c). Notably, three sibling pairs ranked near the bottom due to genetic overlap from close kinship, as expected. Compositional analysis of chromosome 7 showed that individuals of European ancestry carried fractions of non-European ancestry, which contributed to their DI scores and marginally to the total diversity captured in the graph. Conversely, the globally representative HPRC cohort demonstrated a more balanced DI score distribution, with individuals exhibiting higher proportions of African ancestry scoring highest (Fig. 2d). Because the DI score can be calculated directly from variant calls, it provides a practical way to preselect individuals for graph construction when the goal is to maximize genetic diversity.

The ancestry composition of variants in the CF-specific graph was evaluated through direct comparison with the HPRC graph. Single-nucleotide variants (SNVs) were categorized according to the population with the highest allele frequency in gno-mAD [34] (*popmax* ; Fig. 2e). Common SNVs (*popmax >* 10%) were robustly captured in both graphs. Rare SNVs predominantly associated with African and East Asian populations were more effectively represented in the HPRC graph. In contrast, rare SNVs associated with European, South Asian, Middle Eastern, and Ashkenazi Jewish ancestries were better represented in the CF-specific graph. Interestingly, rare SNVs linked to admixed American ancestries were represented in complementary ways, with each graph capturing variants missed by the other.

### 2.2 PangyPlot enables efficient browsing of variation graphs

PangyPlot was developed as a pangenome browser designed to address the limitations of existing visualization tools while remaining scalable to the size of human graphs. Its design is built around three core principles: reliance on a primary reference, bubble-aware visualization, and a force-directed graph layout. First, PangyPlot uses a single reference path that allows regions of interest to be queried directly using standard linear genomic coordinates and enables annotations to be rendered directly on the graph (Fig. 3a). Setup requires a GFA file and the reference path name, with optional gene annotations in GFF3 format to support gene-based search and visualization (setup workflow shown in Supplementary Fig. 9). Second, PangyPlot uses bubbles calculated by BubbleGun [10] to organize complex variation into hierarchical bubbles and chains, which can be interactively expanded to reveal finer detail (Fig. 3b). Finally, PangyPlot relies on an initial two-dimensional layout generated by `odgi layout` [17] but uses a D3 force-directed graph engine [36] to dynamically arrange nodes and links in real time, reducing the need for manual refinement of node positioning (Fig. 3c). The resulting graph is displayed as nodes and links that are strategically rendered to convey relative size differences of the graph components (Fig. 3d).

**Fig. 3.**
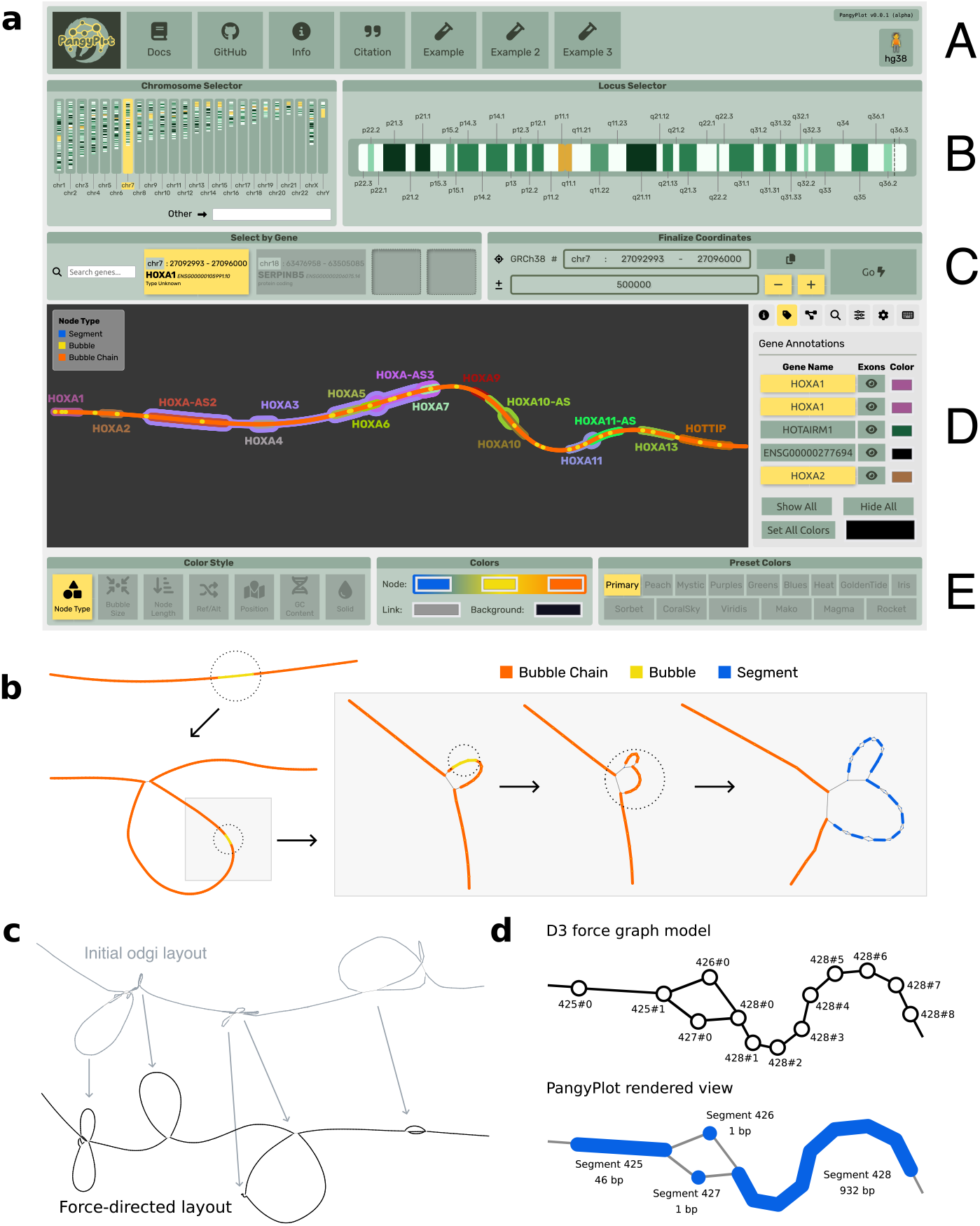
PangyPlot interface for interactive pangenome visualization and core visualization principles. **a**, The PangyPlot user interface showing the *HOXA* gene cluster. (A) Navigation bar preloaded with example regions. (B) A customizable chromosome and locus selector. (C) Genebased search bar (left) and direct coordinate entry (right). (D) The canvas displaying the graph (left) and a control panel with settings and interactive features (right). (E) Color style and palettes. **b**, PangyPlot initially displays top-level bubbles (i.e., those not nested within another bubble; top left, bubble chain in orange, individual bubbles in yellow). When a bubble is “popped” (circle annotation), its internal structure is revealed. Bubbles can be nested, and repeated popping progressively exposes finer levels of variation until the graph resolves to individual segments (blue). **c**, PangyPlot initializes each graph segment with two-dimensional coordinates computed by `odgi layout` (top). These coordinates are subsequently refined in real time using a D3 force-directed physics model, which incorporates node repulsion, collision handling, and link-length optimization to improve the overall layout (bottom). Arrows indicate matching structures. **d**, Rendered view versus D3 implementation. Short segments (e.g., Segment 426, 1 bp) are represented as single D3 nodes. Long segments (e.g., Segment 428, 932 bp) are drawn as thickened polylines but composed of multiple D3 nodes (`428#0–428#8`). This design allows relative segment sizes to be visualized while enabling long segments to bend naturally under the force-directed layout.

To showcase PangyPlot’s functionality, we visualized two CF-relevant loci—one representing small-variant variation and the other structural variation—and compared the results with existing visualization tools. We first examined a 12 kb region of the CF-specific chromosome 7 variation graph that includes exons 10 and 11 of the *CFTR* gene and an upstream polyT repeat that can alter splicing (Fig. 4a). Exon 11 also harbors the most common CF-causing variant (p.Phe508del), consistently observed in *cis* with the (TG)_10_T_9_ allele. PangyPlot setup required approximately 90 minutes to process the full 9 GB chromosome 7 graph, a one-time step that enables repeated queries without reprocessing. The *CFTR* locus was then retrieved by gene name and displayed with integrated annotations (Fig. 4b). For comparison, we visualized the same region with Bandage, Sequence Tube Map, and `odgi`. Both Bandage and Sequence Tube Map required graph conversion to `vg` format and subgraph extraction, which introduced coordinate offsets. After conversion back to GFA, Bandage was capable of a similar visualization to PangyPlot (Fig. 4c) but required custom exon annotations and manual refinement of every node position. In Sequence Tube Map (Fig. 4d), the interweaving lines were difficult to interpret without additional manual annotation, and interpretation diminished further when complex variation was introduced (Supplementary Fig. 10). The `odgi` visualization commands also required conversion to its native format. The two-dimensional `odgi draw` visualization was ineffective at producing images of small variants within this region (Supplementary Fig. 11), but `odgi viz` produced interpretable, linearized images (Fig. 4e). While both Sequence Tube Map and `odgi viz` effectively display haplotype distributions, the resulting visual density obscures small variants. In our case, exons 10 and 11 had to be visualized separately and then combined in an image editor, whereas PangyPlot and Bandage allowed the two exons to be dragged together and viewed simultaneously.

**Fig. 4.**
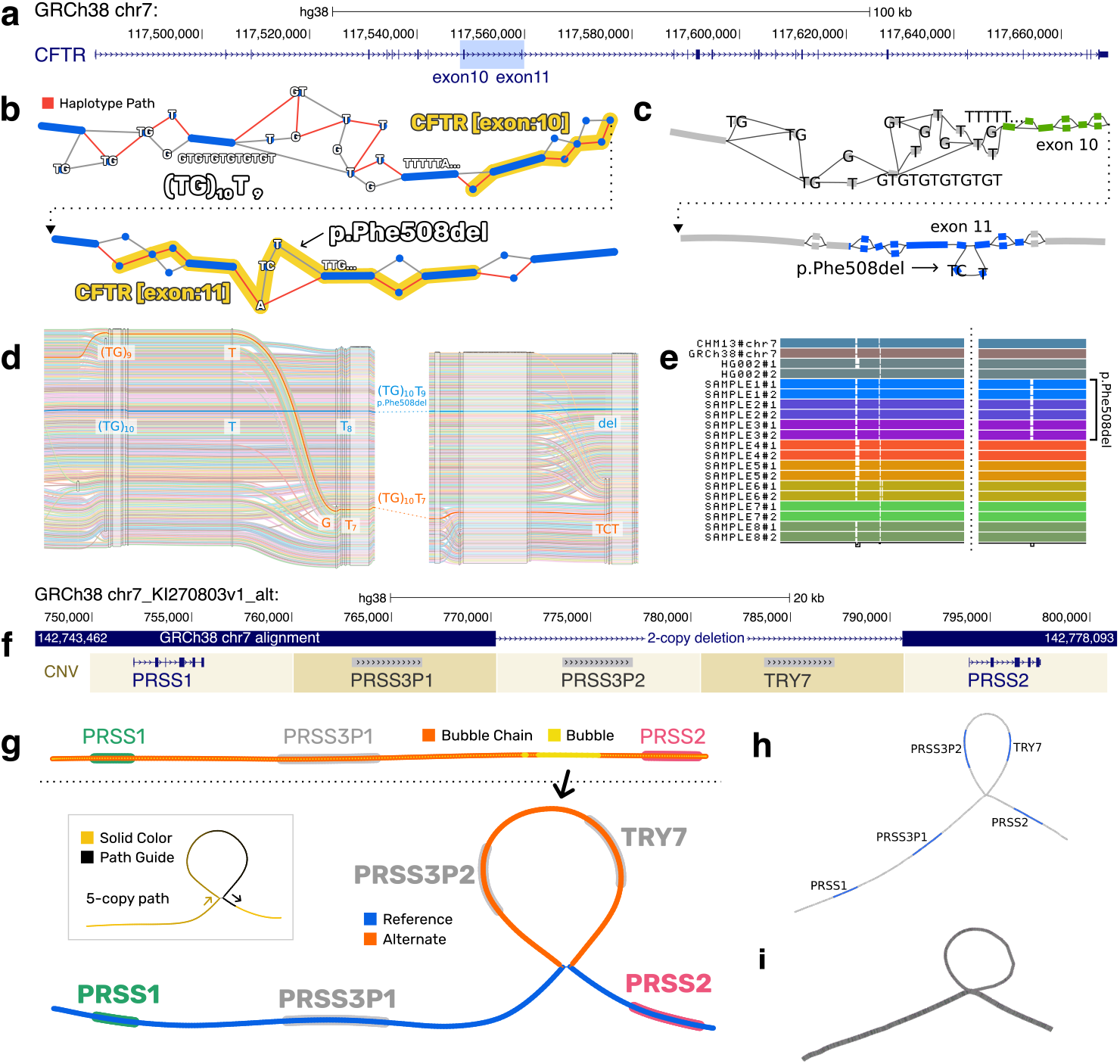
Comparison of graph visualization tools for examining *CFTR* exons 10 (polyT variant), exon 11 (p.Phe508del), and the *PRSS1–PRSS2* region. **a**, Linear view of the *CFTR* gene on GRCh38. Targeted region for visualization is highlighted in blue (exons 10 and 11, chr7:117,548,064–117,560,063). **b**, PangyPlot visualization showing annotated *CFTR* exons (yellow). A single haplotype with (TG)_10_T_9_ and p.Phe508del in *cis* is highlighted via red links. **c**, Bandage visualization with exons 10 and 11 manually annotated. **d**, Sequence Tube Map displays haplotypes as colored lines traversing sequence segments represented by white boxes. Two representative haplotype paths are highlighted: the (TG)_10_T_9_ allele containing the common CF-causing p.Phe508del deletion (blue) and the (TG)_10_T_7_ allele without p.Phe508del (orange). The view was manually truncated to the region of interest and annotations added during post-processing. **e**, A pair of one-dimensional visualizations from `odgi viz`. A representative subset of samples was chosen; haplotypes are drawn as rows. PolyT (left plot) and p.Phe508del (right plot) alleles are shown. **f**, Linear view of the *PRSS1–PRSS2* region on GRCh38 alternative contig KI270803.1, with GRCh38 alignments shown (chr7:142,743,462–142,778,093). KI270803.1 represents the 5-copy allele; the main chr7 contig uses the 3-copy allele. Functional trypsinogen genes are annotated in blue, pseudogenes in grey. **g**, PangyPlot visualization depicting the *PRSS1–PRSS2* region. The initial rendering (top) shows top-level bubbles only. Expanding (“popping”) the large bubble reveals additional CNV copies. The “REF/ALT” color style marks the GRCh38 reference path in blue and alternative segments in orange. The path animation feature (inset) enables real-time viewing of a single path highlighted along the graph. **h**, Bandage visualization of the *PRSS1–PRSS2* region. The BLAST search feature was used to identify trypsinogen genes, and text annotations were added manually. **i**, The `odgi draw` rendering of the *PRSS1–PRSS2* region; line thickness indicates path depth (allele frequency).

We subsequently visualized the CF-associated *PRSS1–PRSS2* trypsinogen locus on chromosome 7 (Fig. 4f; [29]), which harbors a common 10 kb CNV. Most individuals carry either three or five copies, each containing a trypsinogen gene or pseudogene with high sequence similarity (*>*90%). Our PacBio CLR-based assemblies failed to consistently resolve this region due to the size and similarity of the repeated units. Attempts to directly call copy number from PacBio HiFi alignments against either GRCh38 or CHM13 (3-copy alleles) failed for individuals with the 5-copy allele due to reference bias (Supplementary Fig. 12). However, HiFi-based *de novo* assemblies reconstructed the CNV correctly. Evaluation of a 64-sample HiFi-based variation graph clearly delineated samples based on CNV allele in Supplementary Fig. 13, with allele frequencies consistent with prior work [37]. Interactive visualization of this locus in PangyPlot represented the CNV as a single bubble that could be expanded to reveal the full five-copy haplotype structure (Fig. 4g). Using the animated path highlighting feature, CNV allele could be visualized for any sample in the graph. Bandage and `odgi draw` similarly showed the CNV as a loop (Fig. 4h,i) but both lacked annotations. The built-in BLAST search in Bandage was used to locate and mark the trypsinogen genes, which required the user to supply the reference sequence. Supplementary Table 3 provides a comparative summary of the visualization tools assessed.

### 2.3 PangyPlot assists in fine-mapping of a repeat-dense locus

Using graph-based analysis in conjunction with PangyPlot, we analyzed the chromosome 5 region p15.33 that was previously implicated as a modifier of lung function in CF [31, 32]. This subtelomeric region is enriched for VNTRs and short repeats, posing a significant challenge for variant characterization and fine-mapping, especially with short reads. We leveraged graph data from both hybrid assemblies (*n* = 101) and HiFi-based assemblies (*n* = 64). Graphs were constructed for a well-assembled region (chr5:311,504–626,711) marked by two independent genome-wide significant SNPs: rs57221529 and rs72711364. Both SNPs were confirmed as eQTLs for multiple nearby genes in GTEx (Table 2). From the graphs, 234 Adotto-annotated repeats [38] were extracted, of which 60.7% showed minimal length variability (standard deviation *<*1 bp). SNP alleles and repeat sequences were jointly derived from each haplotype path, providing phased calls per assembly. After Bonferroni correction, repeat length was significantly associated with rs72711364 at 43 repeats and with rs57221529 at 23 repeats (Supplementary Data 1; Supplementary Fig. 14).

**Table 2.**
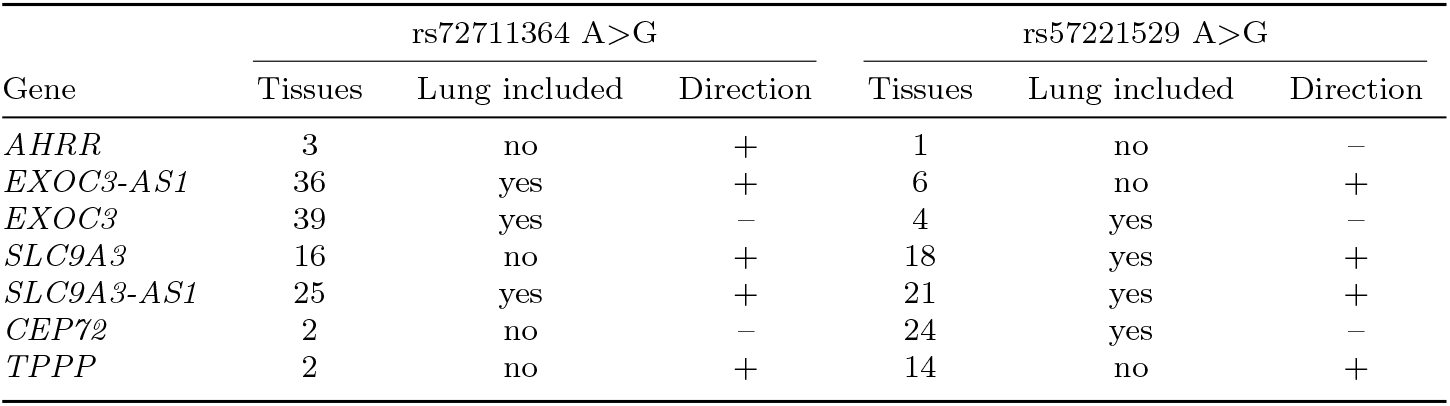
eQTL analysis of lead CF lung function GWAS SNPs in GTEx v10 [40]. Summary of significant eQTL associations for two GWAS SNPs (rs72711364 and rs57221529) previously linked to CF lung function. Shown are the numbers of GTEx tissues with significant eQTL effects, whether the effect is significant in lung tissue, and the expression direction (“+” indicates increased expression associated with the alternative allele G relative to the protective reference allele A). Notably, *EXOC3* and its antisense transcript *EXOC3-AS1* show opposing directional effects in multiple tissues.

| Gene | rs72711364 A>G |  |  | rs57221529 A>G |  |  |
| --- | --- | --- | --- | --- | --- | --- |
|  | Tissues | Lung included | Direction | Tissues | Lung included | Direction |
| *AHRR* | 3 | no | + | 1 | no | – |
| *EXOC3-AS1* | 36 | yes | + | 6 | no | + |
| *EXOC3* | 39 | yes | – | 4 | yes | – |
| *SLC9A3* | 16 | no | + | 18 | yes | + |
| *SLC9A3-AS1* | 25 | yes | + | 21 | yes | + |
| *CEP72* | 2 | no | – | 24 | yes | – |
| *TPPP* | 2 | no | + | 14 | no | + |

For illustrative purposes, we used PangyPlot to follow up on three rs72711364-associated repeats that showed highly polymorphic length distributions (Fig. 5a–c). Fig. 5d shows the relative position and GC content of the three repeats at this locus. Repeat 70 (34 bp VNTR in *AHRR* intron, *p* = 6.24 × 10^*−*16^) overlaps two ENCODE regulatory elements and a H3K27Ac peak and displays large, expanded segments (Fig. 5e). Repeat 87 (polyT in *EXOC3-AS1, p* = 2.70 × 10^*−*10^) occurs within the tail of an AluY element (Fig. 5f). Repeat 88 (GGC trinucleotide in the *EXOC3* 5^*′*^ UTR, *p* = 3.57 × 10^*−*13^), located 46 bp downstream of the transcription start site (TSS), contained 3 to 20 perfect GGC copies. A detailed breakdown of the haplotype distribution (Fig. 5g) and PangyPlot visualization of Repeat 88 (Fig. 5h) is shown. The visualization reveals C*>*G substitutions within the GGC repeat that produce GGGGG homopolymers, forming a sequence pattern strongly consistent with canonical G-quadruplex structures that are known to modulate transcriptional activity [39]. These repeats represent a subset of the significant association results but demonstrate how PangyPlot can be used to better understand the structure of repeats and support hypothesis generation.

**Fig. 5.**
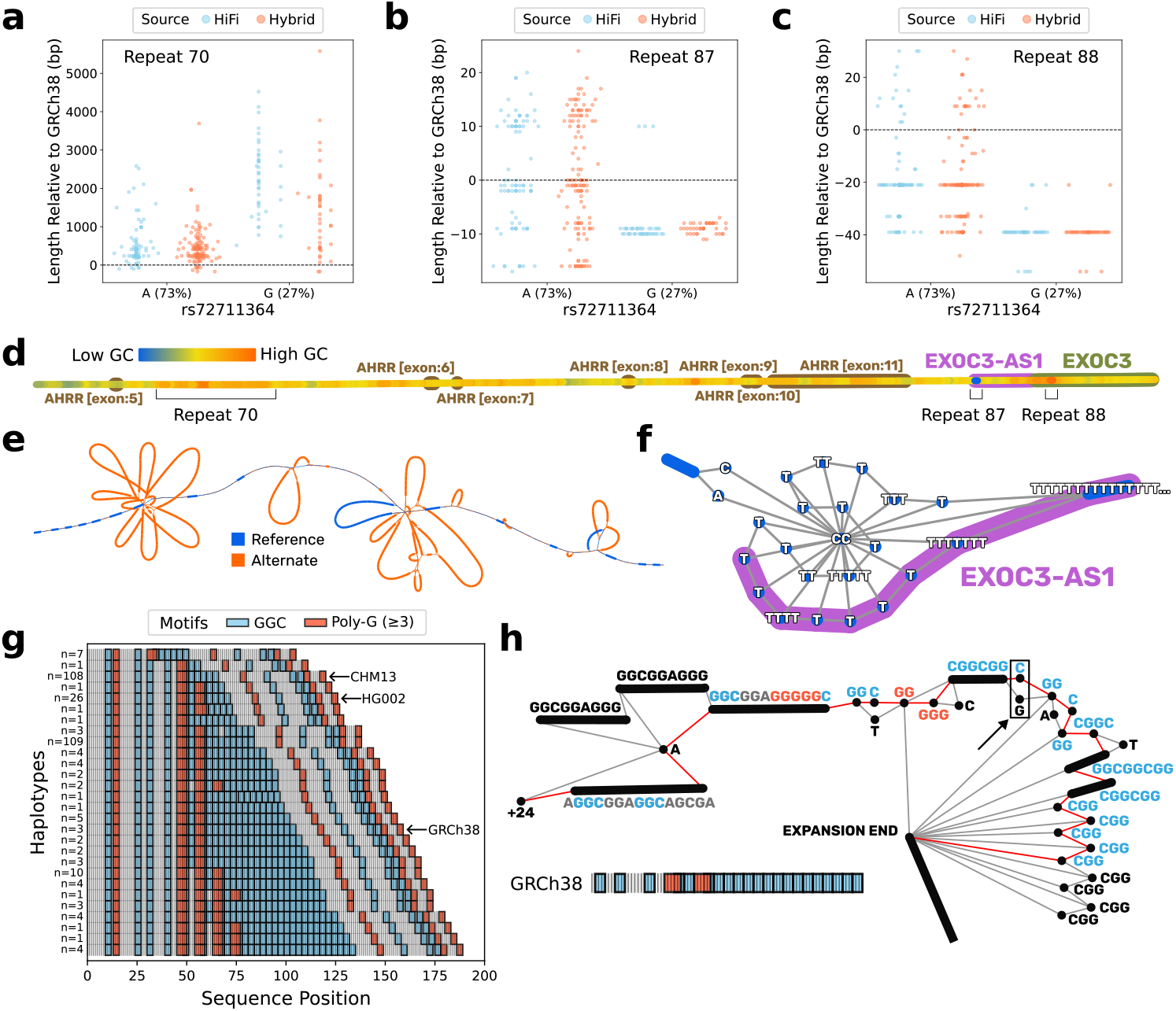
PangyPlot visualization of rs72711364-associated repeat expansions across the *AHRR–EXOC3* region. **a–c**, Repeat length distributions stratified by rs72711364 allele. Each dot represents an individual haplotype, colored by variation graph: blue (HiFi), orange (hybrid). The CF lung function risk allele is G. **d**, PangyPlot visualization showing the relative position of Repeats 70, 87, and 88 (top-level bubbles only). *AHRR* exons (brown), *EXOC3-AS1* (magenta), and *EXOC3* (green) are annotated. Nodes are colored by GC content. Repeat 70 lies in an intron of *AHRR*, Repeat 87 is the low-GC 3^*′*^ end of *EXOC3-AS1*, and Repeat 88 lies in the high-GC 5^*′*^ UTR of *EXOC3*. **e**, Expanded PangyPlot view of Repeat 70 in the `HiFi clip graph`: a 34-mer VNTR (AATCT-GCCTGGTCGGGTGGGAGGCCTAGGGGCCG). Nodes are colored by reference status (blue, on GRCh38 path; orange, alternative relative to GRCh38). **f**, Expanded PangyPlot view of Repeat 87 in the HiFi clip graph: a simple polyT repeat at the 3^*′*^ end of *EXOC3-AS1* terminating on CC. **g**, Linear haplotype distribution of Repeat 88 alleles. Rows represent distinct alleles, colored by sequence motif: GGC repeats (blue) and G-rich homopolymers (≥ 3 bp; orange). The y-axis shows the number of haplotypes carrying each allele. Reference alleles are noted; HG002 is homozygous. **h**, Expanded PangyPlot view of Repeat 88 in the hybrid `d9-filter` graph, beginning +24 bp from the *EXOC3* TSS. Colored segments highlight GGC repeats (blue) and G-rich motifs (orange). The GRCh38 reference haplotype (red path links) is shown, and a black arrow indicates a C*>*G substitution that transforms a pair of GGC repeats into a GGGGG homopolymer.

## 3 Discussion

Public linear reference genomes have been critical in standardizing genomic research, providing shared coordinate systems and comprehensive annotations that enable consistency and comparability across studies. However, despite their benefits and ongoing improvements, linear references inherently represent only a single instance of a genome and therefore exhibit reference bias. This bias can profoundly impact our understanding of genetic diseases and exacerbate health disparities, particularly in populations genetically distant from the reference.

The *PRSS1–PRSS2* locus exemplifies the effect of reference bias. A common CNV in this region induces a strong reference bias effect known to produce clinical misinterpretations with short-read data [41]. This bias persists even with long and accurate PacBio HiFi alignments (Supplementary Fig. 12) across both GRCh38 and CHM13 references. Previous GWAS identified associations between common SNPs at this locus and risk of pancreatitis or CF-related meconium ileus, attributing causal effects to *PRSS1* expression [29, 30]. These studies did not account for the CNV copies, in part because reliance on a single linear reference sequence masks the true extent of variation at the locus. Accounting for reference bias demonstrated that the association is instead likely driven by the CNV itself, with *PRSS2* expression the putative contributing factor [37, 42, 43]. This case illustrates how reference bias can hinder or misinform disease interpretation. In contrast, pangenome-based visualizations make this large CNV immediately apparent, emphasizing the benefit of a variation-aware approach.

Yet, human pangenome references, such as those developed by the HPRC, remain relatively underutilized. Adoption of graph-based approaches has been hindered by both the inherent complexity of graph models and by technical barriers imposed by a field still in the process of establishing shared standards and effective tools. HPRC data are released as multiple graphs constructed using different methods, filters, and formats, reflecting heterogeneity of emerging pangenome standards. Performing graph operations typically requires familiarity with the various GFA specifications in addition to the `vg` [16] and `odgi` [17] toolkits, which together share a combined set of 78 partially overlapping subcommands at the time of writing. Consequently, visualizing specific genomic regions of interest requires significant expertise to get the data into the correct format, and existing visualization tools are generally inflexible in the types of views they can produce and have limited scalability. This stands in contrast to the relative simplicity and flexibility of linear genome browsers.

We developed PangyPlot to lower these technical barriers and help accelerate the adoption of pangenomes. Specifically, PangyPlot can be deployed on a public server and preloaded with graph data, enabling non-expert users to explore pangenomic resources without needing to directly manipulate graph files. An instance preloaded with HPRC data is available at https://pangyplot.research.sickkids.ca, and others can similarly deploy and host PangyPlot instances to accompany their own human or non-human pangenome releases. In addition to being a public utility, PangyPlot can be run locally with custom graph data. To demonstrate this, we generated and visualized multiple variation graphs from a Canadian CF cohort.

Using a hybrid assembly strategy that combined linked-read and long-read sequencing, we constructed a chromosome 7 graph that, after pruning, was comparable in size to the HPRC chromosome 7 graph. Although the CF cohort was predominantly of European ancestry, the graph nevertheless captured substantial ancestral diversity. Rare SNVs associated with European, South Asian, Middle Eastern, and Ashkenazi Jewish ancestries were more effectively represented in the CF graph than in the HPRC graph, despite the latter being explicitly designed to maximize global diversity. Importantly, the CF graph included more than twice as many samples, suggesting that limited ancestral breadth can be partially offset by increased sample depth. The DI score provides a quantitative framework to optimize this balance between ancestral diversity and sample depth when designing future pangenome cohorts.

Visualizing the *CFTR* locus from the 9 GB chromosome 7 GFA file underscored common limitations of existing tools. Interactive browsers such as Sequence Tube Map and Bandage required file conversion and subgraph extraction to accommodate the file size. These steps introduced positional offsets that are appended to path names, complicating queries for specific genomic ranges. Current visualization methods also lack contextual landmarks such as linear reference coordinates and gene annotations, making it difficult to orient within the graph—much like navigating a map without labeled streets. Finally, the generation of publication-quality figures typically requires extensive manual refinement.

PangyPlot was developed to address these three bottlenecks: data preparation, visual orientation, and figure production. While the requirement of a primary linear reference may appear counterintuitive, it provides a familiar framework that simplifies coordinate queries. It also enables gene-based searches and integrated annotations, allowing rapid visual orientation. In addition, abstraction of complex variation into bubble structures and use of a dynamic force-graph layout reduce the effort needed to generate high-quality figures suitable for publication. Rather than replacing existing tools, PangyPlot complements them as a general-purpose browser for targeted regions. Sequence Tube Map excels at small-variant haplotype visualization, `odgi` at large-scale structures, and Bandage at quickly viewing small GFA files.

Furthermore, we demonstrated how hypothesis generation and fine-mapping can be performed with graph-based approaches in conjunction with PangyPlot visualizations. The subtelomeric region on chromosome 5 was previously associated with CF lung function [31, 32], with two independent association signals flanking *SLC9A3*, a sodium/hydrogen exchanger. *SLC9A3* is a strong candidate modifier gene given *CFTR*’s complementary role in ion regulation at the apical membrane, as well as gene-set evidence linking *SLC9A3* to lung disease [44]. However, this associative evidence does not preclude other genes at this locus (*AHRR, EXOC3, CEP72, TPPP*, and antisense transcripts *SLC9A3-AS1* and *EXOC3-AS1*) from being causal.

This region is enriched with highly polymorphic repeats; we identified multiple repeats with length significantly associated with the GWAS SNPs. Among the repeats where length was highly variable and associated with the rs72711364 allele, a highly polymorphic GGC-repeat expansion within the 5^*′*^ untranslated region of *EXOC3* was particularly intriguing. Shorter repeat lengths were associated with the lung function risk allele, and longer repeats contained a set of GGGGG repeats. PangyPlot-enabled visualization of repeat structure showed that these poly-G motifs likely arose from point mutations within the original GGC repeat motif. Such extended guanine-rich regions can facilitate formation of G-quadruplex and R-loop structures [45], potentially altering local transcription dynamics and explaining observed opposing eQTL effects of *EXOC3* and its antisense transcript, *EXOC3-AS1. EXOC3* encodes the Sec6 protein—a component of the exocyst complex critical to vesicular trafficking and secretion. CF pathophysiology is heavily tied to the epithelial membrane landscape (*CFTR* and mucins), and thus the role of *EXOC3* is compelling and warrants further investigation. Importantly, this analysis illustrates how graph-based visualizations can assist in hypothesis generation; further work is necessary to determine whether the repeat expansion in *EXOC3* contributes to lung function or altered gene expression.

In summary, the critical advantage of graph-based references over linear genomes lies in their improved ability to capture, represent, and interpret complex human genetic variation, thereby mitigating the effects of reference bias and enabling improved fine-mapping. PangyPlot substantially enhances accessibility and interpretability of graph-based genomic resources. Nonetheless, PangyPlot’s utility fundamentally depends on underlying graph quality; inaccuracies and assembly errors can introduce unnecessary complexity and impede biological interpretation. Future efforts must also prioritize optimizing graph topologies for biological accuracy and visualization simplicity. Ultimately, broad adoption of standardized, accessible variation graph resources and powerful visualization tools has significant potential to reduce biases in disease research, improve scientific accuracy, and promote more equitable genomic medicine.

## 4 Methods

### DNA sample acquisition

The Canadian CF Gene Modifier Study (CGMS) was approved by the Hospital for Sick Children Research Ethics Board (approval number #1000065760). Blood samples were collected from individuals with CF across Canada and processed at The Hospital for Sick Children (Toronto, Canada).

### 10x Genomics linked-read sequencing

DNA extraction and library preparation have been previously described [46]. Sequencing was performed on the Illumina HiSeq X platform using 150 bp paired-end reads at an average of 30× coverage (∼24× after barcode removal), with an average molecular length of 60 kb. `Long Ranger v2.2.2` [47] was used to align reads to GRCh38 and call variants. *De novo* assembly was performed using `Supernova v2.1.1` [48].

### PacBio continuous long reads and HiFi sequencing

DNA extraction and library preparation have been previously described [46]. Libraries were sequenced on one of three PacBio platforms: Sequel I (34 CLR samples), Sequel II (67 CLR samples), or Revio (64 HiFi samples). Sequel I runs produced reads with an average N50 of 27 kb at 50× coverage; Sequel II runs achieved an average N50 of 29 kb at 76× coverage. HiFi samples had an average read length of 16 kb and 29× coverage. The 101 CLR samples were assembled using `Canu v1.8` [49]. `Hifiasm v0.21.0` [50] was used to assemble HiFi reads, generating two phased haplotype FASTA files per sample.

### Hybrid *de novo* assembly

Full details are provided in Supplementary Method 1. In brief, we employed a multistep approach to assemble and refine 101 phased hybrid *de novo* assemblies using 10XG and PacBio CLR reads. For each sample, a `Supernova` “pseudohap” assembly and a `Canu` assembly were taken as input and refined with `purge dups v1.1.2` [51] (following the official guide for CLR reads) to remove haplotigs. The `Supernova` assembly was split into 1,000 bp chunks and aligned to the `Canu` assembly using `BLASTn v2.7.1` [52]. A custom Python pipeline [53] was then used to stitch the two assemblies into a hybrid assembly (detailed in Supplementary Methods 1). Sequentially aligned and uniquely mapped chunks defined shared regions between the assemblies, facilitating scaffolding, gap filling, and correction of misassemblies. `SDA denovo v1.0.0` [54] was used to manage duplicated sequences.

To phase the assembly, we applied an iterative polishing process using both 10XG and PacBio CLR reads. CLR reads were aligned to the hybrid assembly using `pbmm2 v1.3.0` [55], followed by an initial round of polishing with the Arrow algorithm (`gccp v1.9.0` [56]). 10XG reads were then aligned using `BWA-MEM v0.7.8` [57], followed by a second round of polishing with `Pilon v1.23` [58]. Heterozygous SNPs consistent between `Long Ranger` (10XG) and `Longshot v0.4.1` [59] (CLR) variant calls were used to phase reads into two haplotypes. The iterative alignment and polishing process was repeated with haplotype-specific reads to generate final haploid assemblies.

### Construction of variation graphs with Minigraph-Cactus

We employed the `Minigraph-Cactus` pangenome pipeline v2.7.2 [14] to construct variation graphs from *de novo* assemblies. To assign *de novo* scaffolds to human chromosomes, each assembly was aligned to CHM13 using `minimap2 v2.24` [60] with the `--asm5` flag, outputting in PAF format. Non-aligning scaffold tips were clipped from the assembly.

We constructed a variation graph using chromosome 7 scaffolds from 101 CF-based hybrid assemblies plus HG002, generated from CLR and 10XG reads [61]. Chromosome 7 sequence from CHM13 and GRCh38 (chr7 plus nine alternative contigs) was also included in the graph.

We also constructed graphs for two targeted loci. For the *PRSS1–PRSS2* locus (GRCh38 chr7:142,038,121–143,088,503), we used 64 `hifiasm` assemblies generated from PacBio HiFi reads, along with CHM13, GRCh38, and the overlapping GRCh38 alternative contig `chr7 KI270803v1 alt`. For the chromosome 5 subtelomeric region (GRCh38 chr5:1–626,897), we generated both `hifiasm` (HiFi, *n* = 64) and hybrid (CLR/10XG, *n* = 93) graphs. Hybrid samples that did not span this locus within a single scaffold were excluded. CHM13, GRCh38, and the alternative contig `chr5 KI270793v1 alt` were also included in this graph.

### Ancestry calling with GRAF-pop

We inferred population structure using 10XG genotype data from 564 individuals with CF, each processed using `Long Ranger` to generate phased variant calls. Ancestry-informative variants were extracted using the `ExtractAncSnpsFromVcfGz.pl` script from the `GRAF-pop v1.0` toolkit [35], and population assignments were inferred using the `grafpop` executable and `SaveSamples.pl` script. Genome-wide ancestry distributions were visualized using a custom `R` script adapted for GRAF-pop outputs. This procedure was repeated for HPRC samples using variant calls from the `Minigraph-Cactus` output (`hprc-v1.1-mc-grch38.vcf.gz`) [8]. A similar analysis was also performed using only SNPs from chromosome 7 to estimate ancestry composition specific to that chromosome.

### Calling variants directly from variation graphs

We generated Variant Call Format (VCF) files using the `deconstruct` command of `vg v1.62.0` with the `-a` and `-e` flags, projecting variants onto the GRCh38 path for both HPRC and CF-specific chromosome 7 clipped graphs. Following graph-based variant projection, VCFs were post-processed following the HPRC protocol. Specifically, nested variants and sites with reference allele lengths greater than 100 kb were removed using `vcfbub` [62] (`--max-level 0 --max-ref-length 100000`), and alleles were subsequently realigned using `vcfwave` [63] (`-I 1000`). VCFs were then normalized, decomposed, and left-aligned using `bcftools norm v1.20` [64] with the GRCh38 chromosome 7 reference. Only biallelic SNPs were retained (`bcftools view -v snps`).

Filtered VCFs were annotated using the Ensembl Variant Effect Predictor (`VEP v112.0`) [65], incorporating global and population-specific allele frequencies from gnomAD v4.1 [34]. The following gnomAD population allele frequency annotations were included: African (AFR), Admixed American (AMR), East Asian (EAS), non-Finnish European (NFE), South Asian (SAS), Ashkenazi Jewish (ASJ), and Middle Eastern (MID). The `popmax` annotation, corresponding to the population with the highest population-specific allele frequency, was also included. Annotated variants from both the HPRC and CF cohorts were then filtered to retain only those overlapping high-confidence, non-difficult regions (from GIAB [66], `GRCh38 notinalldifficultregions.bed`). Variants were classified as shared or cohort-specific using `bcftools isec`.

### Diversity Importance (DI) Score

To quantify each sample’s contribution to the genetic diversity represented in the genome graphs, we developed a score based on minor allele frequencies (MAFs). For a given sample, the score is defined as the sum of the inverse MAFs of all alleles present:

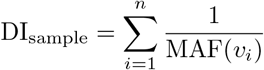

where *n* is the number of variants observed in the sample, *v*_*i*_ represents the *i*th variant, and MAF(*v*_*i*_) is the MAF of that variant within the target population. The DI score captures both the number and rarity of variants in a sample, prioritizing those that introduce uncommon or unique genetic variation into the graph. Samples with more variants naturally contribute more to diversity, and those with rarer variants are weighted even more heavily. To minimize the impact of potential false positives, only biallelic SNPs with non-zero allele frequency in gnomAD v4.1 were included.

### Visualization of *CFTR* with Sequence Tube Map

To visualize the *CFTR* locus using Sequence Tube Map (v0.1.0) [19], we first converted the hybrid chromosome 7 `d9-filter` graph from GFA to `vg` format using `vg convert` (`vg v1.66.0`). We then used `vg chunk` to extract the region spanning GRCh38 chr7:117,361,861–117,786,430, which includes the *CFTR* gene. The resulting subgraph was indexed using `vg index`. To target *CFTR* exons 10 and 11 specifically, we manually created a BED file defining the regions: exon 10, chr7:117,548,595–117,548,662; exon 11, chr7:117,559,556–117,559,642.

Using the `prepare_chunks.sh` script, we generated exon-specific data directories and placed them into the `sequenceTubeMap/exampleData` folder. The Sequence Tube Map server was built and launched locally. The visualization was configured to display GRCh38 paths in black and all other haplotypes in grey; SVG snapshots were exported.

### Visualization of target regions with Bandage

To visualize *CFTR* with Bandage (v2025.5.1) [18], we extracted the region of interest using the same extraction protocol as for Sequence Tube Map, then reconverted it back to GFA format with `vg convert`. The resulting GFA was loaded directly into Bandage. Within Bandage, we created a BLAST database from the graph and queried it with exon 10 and 11 sequences of *CFTR* and the full *PRSS1* gene for the *PRSS1– PRSS2* region. Matching nodes were highlighted based on BLAST alignment, and the node layout was manually adjusted to produce publication-quality SVG figures.

### Visualization of target regions with odgi

To visualize with `odgi` (v0.9.2) [17], GFA graphs with `W` lines were first converted to `vg` format and back to GFA with `P` lines using `vg convert (--no-wline)` for compatibility with `odgi`. GFA files were then converted to `odgi` format using `odgi build`, and the target regions were isolated with `odgi extract`. For 1D visualizations, the graph was sorted with `odgi sort` and rendered using `odgi viz`. We specified relevant haplotypes using `--paths-to-display` and zoomed in on exon regions with `--path-range`, maintaining single-base resolution via `--bin-width=1`. For 2D visualizations, the sorted graph was laid out using `odgi layout` and rendered with `odgi draw`, optimizing layout parameters for clarity.

### Visualization of target regions with PangyPlot

To visualize with PangyPlot, GFA files were converted to `odgi` format following the same conversion protocol described above. The graph was sorted in one dimension (`odgi sort --optimize --Y`, with the `--H` parameter to prioritize the GRCh38 path). Two-dimensional layout coordinates were computed with `odgi layout (--tsv)`. A GFA file was produced using `odgi view`. Data were indexed using `pangyplot add`, given the GFA and layout file. GFF3 annotation files were downloaded from GENCODE (Release 48) [67] and indexed using `pangyplot annotate`. For HPRC data, individual chromosomes from the “`hprc-v1.1-mc-grch38`” `clip` graphs were downloaded, and this process was followed to generate the PangyPlot instance available at https://pangyplot.research.sickkids.ca.

### Analysis of repeat structures in chromosome 5 and association with GWAS SNPs

Sequence segments corresponding to SNPs rs57221529 and rs72711364 were extracted from the chromosome 5 graphs. Variants were manually genotyped as A or G based on the segment traversed for each haplotype path. Within the target region (chr5:311,504–626,711), 234 repeats were cataloged using Adotto v1.2.1 [38], encompassing both short tandem repeats and VNTRs. A custom script was written to extract local subgraphs around each repeat using `odgi extract`, and to convert paths within each subgraph to FASTA format using `odgi paths (-f)`. This produced haplotype-resolved FASTA sequences for each repeat.

To assess the relationship between repeat length and SNP alleles, we computed the repeat length for each haplotype and performed logistic regression using the SNP allele as the outcome variable. Covariates included assembly type (HiFi or hybrid) and sex. Summary statistics (mean, standard deviation, and range) were computed separately for each allele group (A vs. G), and counts of unique repeat lengths were recorded. Visualizations and association test outputs were generated per repeat–SNP pair.

## Supplementary information

Supplementary Information is available for this paper. It includes Supplementary Methods, Supplementary Figures 1–14, and Supplementary Tables 1–3. An accompanying Excel file (`Supplementary Data 1.xlsx`) contains repeat association statistics.

## Acknowledgements

We thank the patients, care providers, clinic research assistants, collaborators, and principal investigators at CF Centers across Canada for their contributions to the Canadian CF Gene Modifier Study (https://github.com/strug-hub/Canadian-Cystic-Fibrosis-Gene-Modifier-Study), the CCF Canada Patient Registry, and the CF Canada–SickKids Program in Individual Therapy.

Funding was provided by the CF Canada 2022 Basic and Clinical Research Grant (1009794), jointly supported by Cystic Fibrosis Canada and the CIHR Institute of Circulatory and Respiratory Health (FRN: BCG 187014); Cystic Fibrosis Canada (Grant 608828); the Cystic Fibrosis Foundation (STRUG17PO); the Canadian Institutes of Health Research (Foundation Grant FRN-167282); and the Government of Canada through Genome Canada and the Ontario Genomics Institute (OGI-148). This research was undertaken, in part, thanks to support from the Canada Research Chairs Program to L.J. Strug, Canada Research Chair in Genome Data Science.

PangyPlot builds upon existing graph algorithms, notably the layout algorithm from `odgi` [20] and the bubble detection algorithm from `BubbleGun` [10]. PangyPlot also extends the D3 force graph implementation by Vasco Asturiano [36]. We thank these developers for making their tools publicly available. We also thank Laurie Lintott for helpful comments on the manuscript draft.

## Declarations

### Authors’ contributions

**Conceptualization:** S.M. and L.J.S.

**Supervision:** L.J.S.

**Funding acquisition:** L.J.S.

**Resources:** K.K., J.A., P.E., F.R., and CCFGMS

**Sample processing:** F.L. and K.K.

**Data curation:** F.L. and K.K.

**Software:** S.M.

**Formal analysis:** S.M.

**Data processing:** S.M., B.T., Z.W., R.V.H., W.W.L.S., and A.H.

**Methodology:** S.M., S.C., D.R., B.T., Z.W., A.H., C.W., and L.J.S.

**Writing:** S.M., with input from all authors.

## Data availability

A live PangyPlot instance preloaded with HPRC data is available at https://pangyplot.research.sickkids.ca.

Hybrid assembly for HG002: https://zenodo.org/records/15062708 HPRC `minigraph-cactus clip` GFA and layout files: https://zenodo.org/records/17173731

PangyPlot database files: https://zenodo.org/records/17174109

## Code availability

The PangyPlot source code is openly available at https://github.com/scottmastro/pangyplot under the MIT License. Detailed usage instructions, API documentation, and example datasets are provided at https://pangyplot.readthedocs.io/. The hybrid assembly pipeline used in this study is openly available at https://github.com/scottmastro/hybrid-pipeline under the MIT License.

## Ethics approval and consent to participate

The Canadian CF Gene Modifier Study (CGMS) was approved by the Research Ethics Board of The Hospital for Sick Children (#0020020214 from 2002–2019 and #1000065760 from 2019 to the present), as well as by all participating sub-sites. Written informed consent was obtained from all participants or their parents/guardians/substitute decision makers prior to inclusion in the study.

## Canadian Cystic Fibrosis Gene Modifier Consortium

Lisa J. Strug^6^, Katherine Keenan^6^, Felix Ratjen^6^, Johanna Rommens^6^, Melinda Solomon^6^, Candice Bjornson^7^, Mark Chilvers^8^, April Price^9^, Michael Parkins^10^, Michael Derynck^11^, Emmanuelle Brochiero^12^, Lara Bilodeau^13^, Dimas MateosCorral^14^, Daniel Hughes^14^, Mary Jane Smith^15^, Nancy Morrison^16^, Janna Brusky^17^, Anne Stephenson^18^, Elizabeth Tullis^18^, Bradey Quon^19^, Jaled Yehya^20^, Winnie M. Leung^20^, Andre Cantin^21^, Larry Lands^22^

^6^The Hospital for Sick Children, Toronto, ON, Canada.

^7^Alberta Children’s Hospital, Calgary, AB, Canada.

^8^BC Children’s Hospital, Vancouver, BC, Canada.

^9^Children’s Hospital of Western Ontario (London Health Sciences Centre), London, ON, Canada.

^10^Foothills Medical Centre, Calgary, AB, Canada.

^11^Kingston Health Sciences Centre, Kingston, ON, Canada.

^12^Centre de recherche du Centre hospitalier de l’Université de Montréal (CRCHUM), Montréal, QC, Canada.

^13^Institut universitaire de cardiologie et de pneumologie de Québec (IUCPQ), Québec, QC, Canada.

^14^IWK Health Centre, Halifax, NS, Canada.

^15^Janeway Children’s Health & Rehabilitation Centre, St. John’s, NL, Canada.

^16^Queen Elizabeth II Health Sciences Centre, Halifax, NS, Canada.

^17^Royal University Hospital, Saskatoon, SK, Canada.

^18^St. Michael’s Hospital, Toronto, ON, Canada.

^19^St. Paul’s Hospital, Vancouver, BC, Canada.

^20^University of Alberta Hospital, Edmonton, AB, Canada.

^21^Centre hospitalier universitaire de Sherbrooke (CHUS), Sherbrooke, QC, Canada.

^22^Montreal Children’s Hospital, Montréal, QC, Canada.

