## Supplemental for "Beyond reference bias: Making pangenomes accessible with PangyPlot"

### Supplementary Information for: ***Beyond reference bias: Making pangenomes accessible with PangyPlot***

### 1 Supplementary Methods

#### Supplementary Method 1: Construction of the phased hybrid assembly

We utilized a multi-step approach to assemble and refine phased hybrid *de novo* assemblies using PacBio continuous long reads (CLR) and 10X Genomics linked reads (10XG) sequenced from the same sample. We applied this procedure to 101 samples from the Canadian Cystic Fibrosis (CF) Gene Modifier Study and the Genome in a Bottle (GIAB) sample HG002. For each sample, two read-specific input haploid assemblies were generated: Canu 1.8 [1] was used for CLR and Supernova 2.1.1 [2] for 10XG ("pseudohap" assembly). Input assemblies were refined using `purge_dups` 1.1.2 [3] (following the official guide for CLR reads) to remove haplotigs. A custom Python package [4] was developed to merge the two assemblies, summarized in Supplementary Fig. 1 and described below.

##### Creating haploid hybrid assemblies using Canu and Supernova as input

The Supernova assembly was fragmented into 1000 bp chunks and aligned against the Canu assembly using BLASTn 2.7.1 [5] (`-evalue 1e-35 -max_target_seqs 10 -max_hsps 5`). From these alignments, blocks of homology were defined where sequential chunks (minimum of three) uniquely aligned and maintained a consistent order and orientation. Larger "superblocks" were similarly established as a series of homologous blocks generally located within the same genomic locus in both assemblies. This allowed for scaffolding, gap filling, and misassembly correction between blocks.

We strategically leveraged the strengths of both technologies. Supernova utilizes DNA barcodes to group reads derived from the same physical molecule of DNA, reducing the likelihood of cross-chromosome misassemblies. Conversely, local regions flanked by low-complexity DNA often caused inversions in the Supernova sequence, which would not occur with Canu because the long reads span across the low-complexity patches. When the assemblies disagreed, we relied on Canu for resolving localized issues (i.e., at the block level) and Supernova for large-scale structural accuracy (i.e., at the superblock level). Additionally, we favored Supernova's higher base-pair accuracy for most of the genome, switching primarily to fill in low-complexity gaps (Supplementary Fig. 2). We also extended the hybrid assembly wherever possible when a Supernova contig extended past a Canu contig or vice versa (Supplementary Fig. 3).

The task of stitching together the two input assemblies into a unified hybrid requires identifying *forks*: unambiguous base-pair coordinates shared by both assemblies that define possible jump points between the two assemblies. These forks were determined through pairwise sequence alignment within a shared block; a list of forks defines a path across both assemblies. A final step connected paths across multiple contigs to form scaffolds, which optionally relied on Canu unitigs to improve confidence. The result is a single unphased hybrid assembly. On average, Supernova contributed 93% of the hybrid sequence content (median segment length: 31 kb), while Canu contributed the remaining 7% (median segment length: 600 bp).

#### Phasing haploid hybrid assemblies with CLR and 10XG reads

To phase the haploid assembly, an iterative polishing approach was employed, summarized in Supplementary Fig. 4. First, the PacBio CLR reads were aligned to the hybrid assembly using `pbmm2` 1.3.0 [6], and 10XG reads were aligned using Long Ranger 2.2.2 [7]. Segmental Duplication Assembler (`SDA denovo`) 1.0.0 [8] was used in combination with the `pbmm2` alignments to identify and split scaffolds that represented large segmental duplications. CLR reads were used to polish the assembly with the Arrow algorithm (gcpp 1.9.0) [9], and 10XG reads were realigned to the polished output with BWA-MEM 0.7.8 [10]. Pilon 1.23 [11] was then used to polish the assembly. This produced a consensus assembly representing an intermediate of both haplotypes.

Next, the PacBio and 10XG reads were realigned using pbmm2 and Long Ranger, respectively. SNPs were called from the CLR data using Longshot 0.4.1 [12], while Long Ranger produced a phased set of 10XG variant calls. A VCF was generated containing the heterozygous variants present in both callsets (i.e., high-confidence heterozygous SNPs). WhatsHap [13] used this VCF to phase the CLR and 10XG reads into two haplotypes; unphased reads were randomly assigned to a haplotype. Finally, using haplotype-specific reads, the same multi-step polishing procedure was repeated twice, resulting in two assemblies corresponding to the two distinct haplotypes. Supplementary Fig. 5 demonstrates how the assembly converges into two distinct haplotypes.

#### Benchmarking hybrid assembly accuracy

We evaluated genome-wide contiguity of the haploid assemblies for all 101 hybrid samples using QUAST 5.0.0 (--eukaryote) [14]. Each assembly was compared to the T2T-CHM13 v2.0 reference genome [15], and summary statistics were averaged across samples (Supplementary Table 1). The hybrid assemblies demonstrated the highest contiguity, averaging 546 scaffolds (N50: 35 Mb, L50: 30) compared to 6,790 scaffolds (N50: 27 Mb, L50: 34) in Supernova and 876 scaffolds (N50: 27 Mb, L50: 38) in Canu. The hybrid approach also had 62% fewer misassemblies than Supernova and 45% fewer than Canu. Canu had the highest indel rates, which were not reflected in the hybrid assembly. Supernova assemblies contained many gaps, but 98.8% of these were closed in the hybrid assemblies, demonstrating that the hybrid approach compensates for weaknesses of the two input assemblies.

To visualize scaffold breaks, haploid chromosome 7 assemblies were aligned to the T2T-CHM13 v2.0 reference using minimap2 2.24 [16] and visualized as blocks alongside segmental duplications from SEDEF [17]. On chromosome 7, most assembly breaks corresponded to large (>30 kb) segmental duplications or the centromere, which cannot be resolved with either 10XG or CLR data.

We applied the phasing step to create diploid chromosome 7 assemblies (average: 48 scaffolds per sample) and for the chromosome 5 subtelomeric region (GRCh38 chr5:1–626,897; resolved in one scaffold for most samples). Phased variant calls were generated using `dipcall` 0.3 [18] and compared to Genome in a Bottle (GIAB) v4.2.1 high-confidence benchmark calls [19]. Benchmarking was performed using `vcfdist` 2.5.3 [20]. For the hybrid HG002 assembly, we observed high recall for both SNPs (99.7%) and indels (96.1%), but lower precision (SNP: 97.7%, indel: 79.4%) suggested an excess of false positives. Most false positives were found in homopolymers or dinucleotide repeats, or within segmental duplications. Benchmarking was repeated across stratified subsets excluding homopolymers (≥4 bp), dinucleotide repeats (≥10 bp), and segmental duplications (>10 kb), using GIAB genome stratification (v3.4) BED files. Excluding these regions increased SNP precision to 99.6% and indel precision to 92.2%.

We compared these results by similarly benchmarking a HiFi + Hi-C assembly of HG002 [21] against the GIAB callset. When excluding problematic repeat regions, both assemblies exhibited comparable F1 scores ($F_1 = 2 \times \frac{\text{precision}\times\text{recall}}{\text{precision}+\text{recall}}$) for SNPs (hybrid: 0.998, HiFi: 0.999) and indels (hybrid: 0.987, HiFi: 0.991). Detailed statistics are shown in Supplementary Table 1. Notably, the hybrid assembly showed fewer phasing errors, consistent with CLR and 10XG reads having improved phasing potential (Supplementary Fig. 7). These results provide confidence in the accuracy of our assemblies for use in variation graph construction.

#### 2 Supplementary Figures

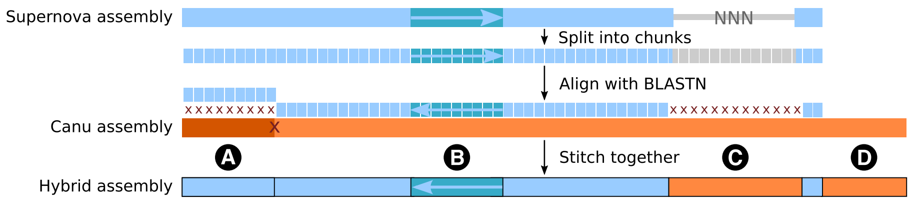


**Supplementary Fig. 1.** Combining Canu and Supernova assemblies into a hybrid assembly. The Supernova assembly was split into chunks and aligned against the Canu assembly using BLASTn. Larger blocks representing the same region were defined and used for error correction. For regions where the assemblies disagreed, one assembly was prioritized over the other based on the strength of each technology. **A**, Misassembly in Canu; Supernova prioritized. **B**, An inversion in Supernova corrected with Canu. **C**, A gap in Supernova filled by Canu. **D**, Extending and scaffolding contigs.

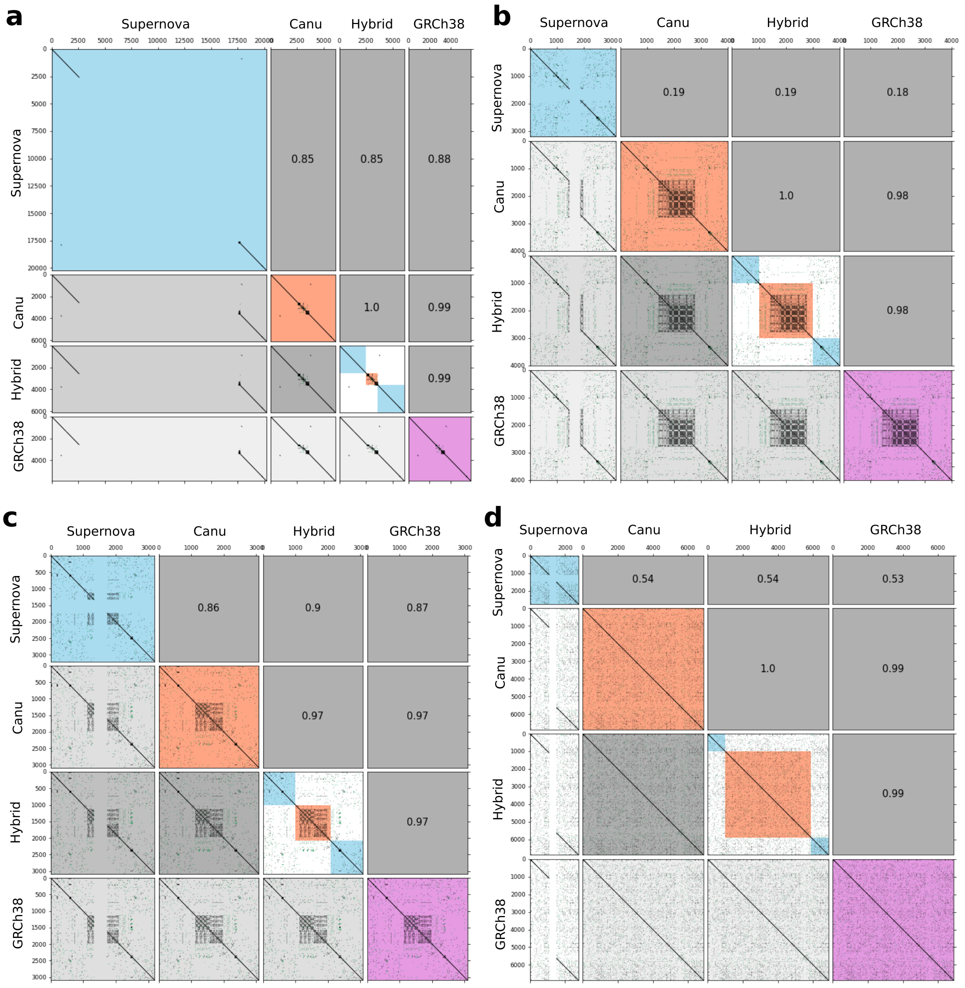


**Supplementary Fig. 2.** Hybrid assembly gap filling of Supernova using Canu. Dot plots for regions with a gap in the Supernova sequence. Gaps are represented as poly-N stretches with lengths estimated by Supernova. Rows correspond to Supernova sequence (blue), matching region identified in Canu (orange), hybrid assembly sequence, and the corresponding sequence from GRCh38 (purple). The upper triangle of each matrix shows the percent sequence identity. **a**, A large 15 kb gap in Supernova replaced by a smaller 1.2 kb sequence from Canu with 99% similarity to GRCh38. **b**, A 1 kb repetitive region filled by Canu. **c**, A partially assembled repetitive region filled. **d**, A case where Supernova underestimates the size of the gap.

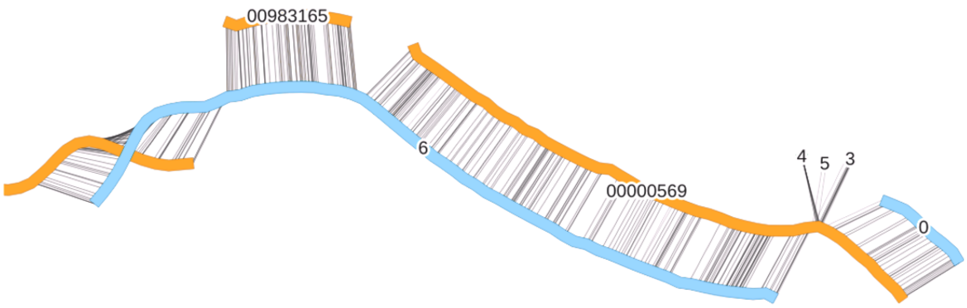


**Supplementary Fig. 3.** Hybrid assembly scaffolding between input contigs.

Scaffolding between Canu (orange) and Supernova (blue); contigs are labeled by their ID, and black lines indicate forks of shared sequence between the assemblies. Scaffolding is performed by jumping between assemblies. For example, Supernova contig 0 can jump to Canu contig 00000569, which can then jump back to Supernova contig 6. In this case, five Supernova contigs and three Canu contigs were scaffolded into a single hybrid scaffold.

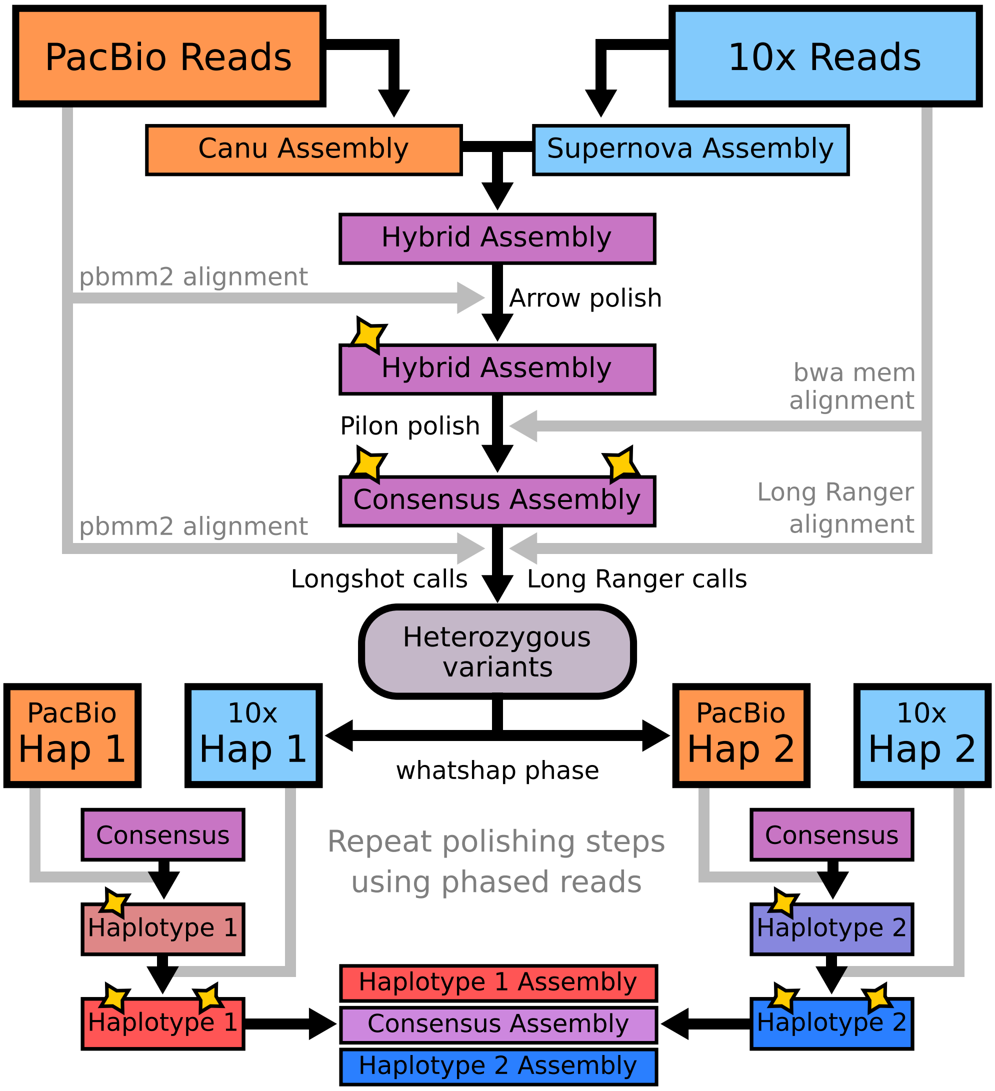


**Supplementary Fig. 4.** Summary of phasing pipeline to produce a single, phased *de novo* genome assembly. DNA from a single individual is extracted and sequenced using PacBio CLR and 10XG technologies. Two assemblies are constructed corresponding to each sequencing technology. The assemblies are merged into a single hybrid sequence and phased using the combined read data from both technologies.

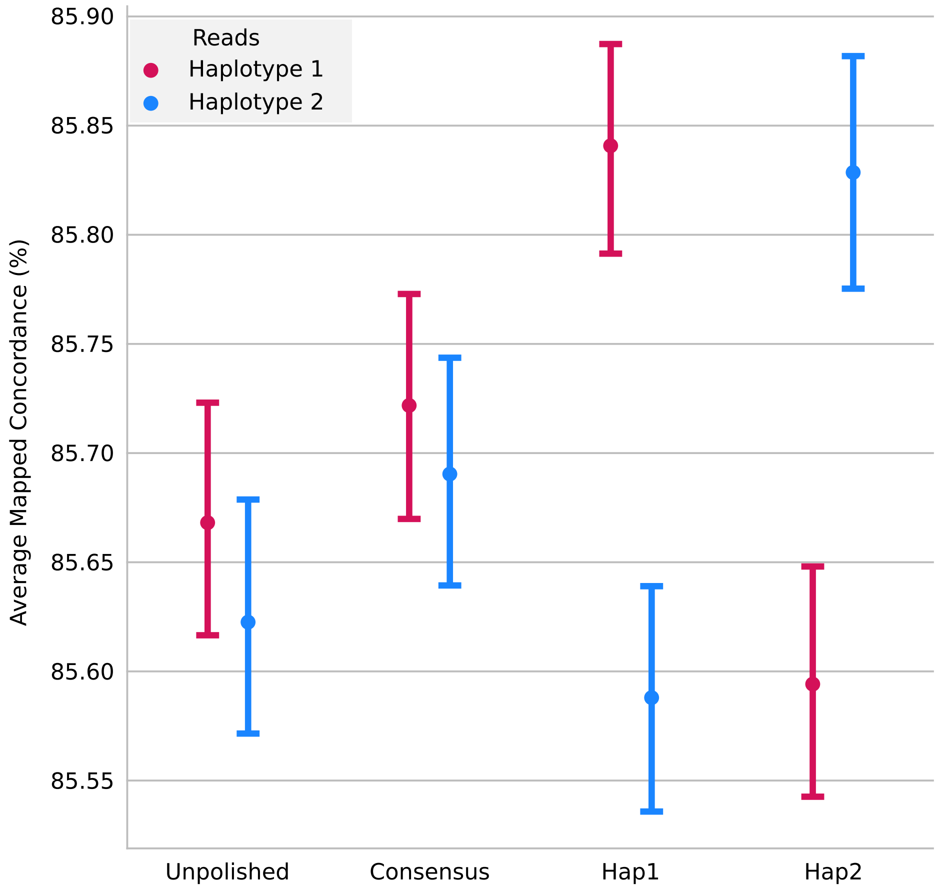


**Supplementary Fig. 5.** PacBio CLR read concordance during haplotype polishing.

To generate two phased haploid assemblies, PacBio CLR and 10XG reads were used to create a polished consensus assembly. This consensus was then polished separately with haplotype-specific reads to generate haplotype 1 and haplotype 2 assemblies. The plot shows average mapped read concordance (%) from `pbmm2` alignments with standard error for a single representative scaffold; reads from haplotype 1 are shown in red and haplotype 2 in blue. Initial consensus polishing improves concordance for both haplotypes, while subsequent haplotype-specific polishing leads to a divergence in concordance values between haplotype 1 and haplotype 2 read groups, as expected from successful phasing. The CLR reads are error-prone, which establishes the upper limit of concordance.

**Supplementary Table 1.** QUAST assembly statistics demonstrate the strength of the hybrid approach.
Haploid genome assembly quality was assessed by QUAST against the reference genome CHM13 ($n$=101). Three assembly methods were compared: Supernova (10XG reads), Canu (CLR), and a hybrid assembly combining both. GRCh38 is provided as a comparison. Scaffold metrics include gaps, while contig metrics break the assembly at gaps before assessment.

| | Supernova (10XG) | | Canu (PacBio) | | Hybrid | | GRCh38 | |
|---|---|---|---|---|---|---|---|---|
| **Type** | Scaffold | Contig | Scaffold | Contig | Scaffold | Contig | Scaffold | Contig |
| Largest contig (Mb) | 98.8 | 2.6 | 97.6 | 97.6 | 104.4 | 64.6 | 249.0 | 141.4 |
| Total length (Gb) | 2.81 | 2.75 | 2.85 | 2.85 | 2.82 | 2.82 | 3.11 | 2.94 |
| GC (%) | 40.84 | 40.84 | 40.89 | 40.89 | 40.90 | 40.90 | 40.88 | 40.88 |
| # contigs | 6790 | 51964 | 876 | 898 | 546 | 913 | 126 | 3279 |
| N50 (Mb) | 27.1 | 0.6 | 27.4 | 27.3 | 35.0 | 15.3 | 145.1 | 57.9 |
| N75 (Mb) | 12.1 | 0.3 | 12.3 | 12.2 | 16.6 | 7.5 | 114.4 | 26.1 |
| L50 | 34 | 6480 | 38 | 38 | 30 | 61 | 9 | 18 |
| L75 | 77 | 14478 | 87 | 87 | 71 | 142 | 14 | 37 |
| # misassemblies | 1213 | 613 | 831 | 810 | 456 | 397 | 7534 | 4745 |
| # local misassemblies | 6552 | 1437 | 2735 | 2731 | 2647 | 2489 | 3894 | 3668 |
| Unaligned length (Mb) | 2.6 | 2.2 | 12.8 | 12.8 | 3.4 | 3.3 | 18.1 | 17.0 |
| Genome fraction (%) | 88.3 | 88.0 | 90.8 | 90.8 | 90.2 | 90.2 | 92.6 | 92.6 |
| Duplication ratio | 1.02 | 1.00 | 1.00 | 1.00 | 1.00 | 1.00 | 1.07 | 1.01 |
| # N's per 100 kb | 2216 | 0 | 0 | 0 | 98 | 0 | 5460 | 0 |
| # mismatches per 100 kb | 102 | 99 | 99 | 99 | 104 | 104 | 126 | 126 |
| # indels per 100 kb | 24 | 22 | 34 | 34 | 26 | 26 | 25 | 25 |
| Largest alignment (Mb) | 35.7 | 1.5 | 77.0 | 77.0 | 81.5 | 54.9 | 93.7 | 93.7 |
| Total aligned length (Gb) | 2.76 | 2.74 | 2.84 | 2.84 | 2.81 | 2.81 | 2.91 | 2.92 |
| NA50 (Mb) | 7.2 | 0.2 | 18.3 | 18.3 | 21.4 | 12.3 | 26.9 | 25.8 |
| NA75 (Mb) | 3.1 | 0.1 | 8.1 | 8.1 | 10.0 | 5.7 | 10.2 | 11.3 |
| LA50 | 114 | 6512 | 52 | 52 | 45 | 75 | 35 | 35 |
| LA75 | 266 | 14563 | 120 | 120 | 106 | 173 | 82 | 78 |

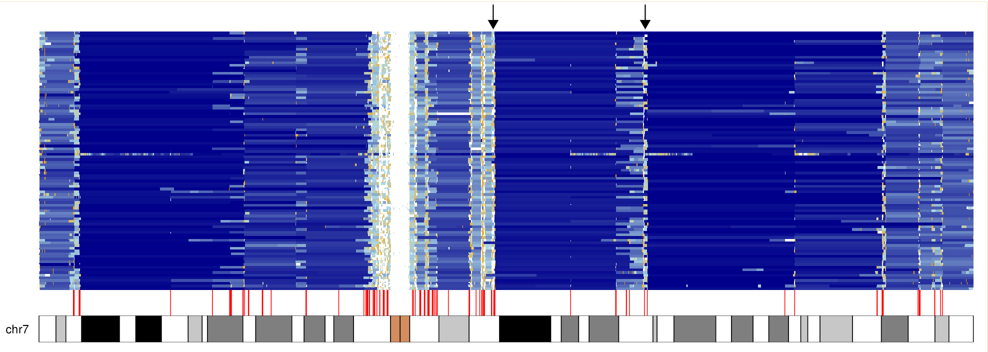


**Supplementary Fig. 6.** Segmental duplications overlap with scaffold breaks.

The hybrid assembled haplotypes were aligned to chromosome 7 of CHM13. Each row represents a haploid hybrid assembly. Scaffolds are colored from longest to shortest on a blue–orange gradient. Segmental duplications identified using SEDEF [17] were filtered to duplications longer than 30 kb and annotated as red lines. Large segmental duplications strongly overlap with assembly breakpoints. Two black arrows indicate the location of a 113 kb duplication with 97% sequence identity. This duplication cannot be resolved with PacBio CLR reads and causes two breaks in all assemblies.

**Supplementary Table 2.** Benchmarking across stratified regions for hybrid HG002 assembly.

Variant calling metrics (TP, FP, FN, precision, recall, F1) and phasing errors (switch, flip, switch/flip NGC50) are shown for SNPs and indels between the Hybrid (PacBio CLR + 10XG) and HiFi (PacBio HiFi + Hi-C) assemblies of HG002 [21]. Stratifications considered: (1) all overlapping regions between the two assemblies, (2) excluding homopolymer (HP) and dinucleotide (DN) repeats, (3) excluding HP/DN and segmental duplications (SD), and (4) excluding all difficult regions. CLR-based hybrid assemblies show reduced precision in indels due to homopolymer errors, while HiFi assemblies perform consistently well across stratifications. Phasing is evaluated using switch and flip error rates.

**SNPs**

| | All overlapping | | No HP/DN repeats | | No HP/DN, no SD | | No difficult regions | |
|---|---|---|---|---|---|---|---|---|
| **Metric** | Hybrid | HiFi | Hybrid | HiFi | Hybrid | HiFi | Hybrid | HiFi |
| TP | 253519 | 255464 | 183624 | 185135 | 173716 | 172724 | 185414 | 184243 |
| FN | 726 | 360 | 480 | 236 | 112 | 169 | 73 | 185 |
| FP | 5957 | 903 | 4080 | 475 | 750 | 248 | 571 | 182 |
| Precision | 0.977 | 0.9965 | 0.9783 | 0.9974 | 0.9957 | 0.9986 | 0.9969 | 0.9990 |
| Recall | 0.9971 | 0.9986 | 0.9974 | 0.9987 | 0.9994 | 0.9990 | 0.9996 | 0.9990 |
| F1 | 0.9870 | 0.9975 | 0.9877 | 0.9981 | 0.9975 | 0.9988 | 0.9983 | 0.9990 |
| Switch Errors | 154 | 363 | 145 | 320 | 132 | 299 | 132 | 314 |
| Flip Errors | 44 | 38 | 17 | 46 | 11 | 47 | 7 | 39 |
| NGC50 (Mb) | 1.4 | 0.7 | 1.7 | 0.7 | 1.9 | 0.8 | 1.6 | 0.8 |

**Indels**

| | All overlapping | | No HP/DN repeats | | No HP/DN, no SD | | No difficult regions | |
|---|---|---|---|---|---|---|---|---|
| **Metric** | Hybrid | HiFi | Hybrid | HiFi | Hybrid | HiFi | Hybrid | HiFi |
| TP | 36518 | 37492 | 10725 | 10717 | 10191 | 10045 | 10955 | 10812 |
| FN | 1450 | 463 | 99 | 103 | 71 | 99 | 9 | 36 |
| FP | 9446 | 4434 | 1590 | 540 | 852 | 511 | 270 | 148 |
| Precision | 0.7942 | 0.8941 | 0.8706 | 0.9519 | 0.9227 | 0.951 | 0.9759 | 0.9865 |
| Recall | 0.9610 | 0.9878 | 0.9909 | 0.9904 | 0.9931 | 0.9902 | 0.9992 | 0.9967 |
| F1 | 0.8700 | 0.9386 | 0.9269 | 0.9708 | 0.9566 | 0.9705 | 0.9874 | 0.9916 |

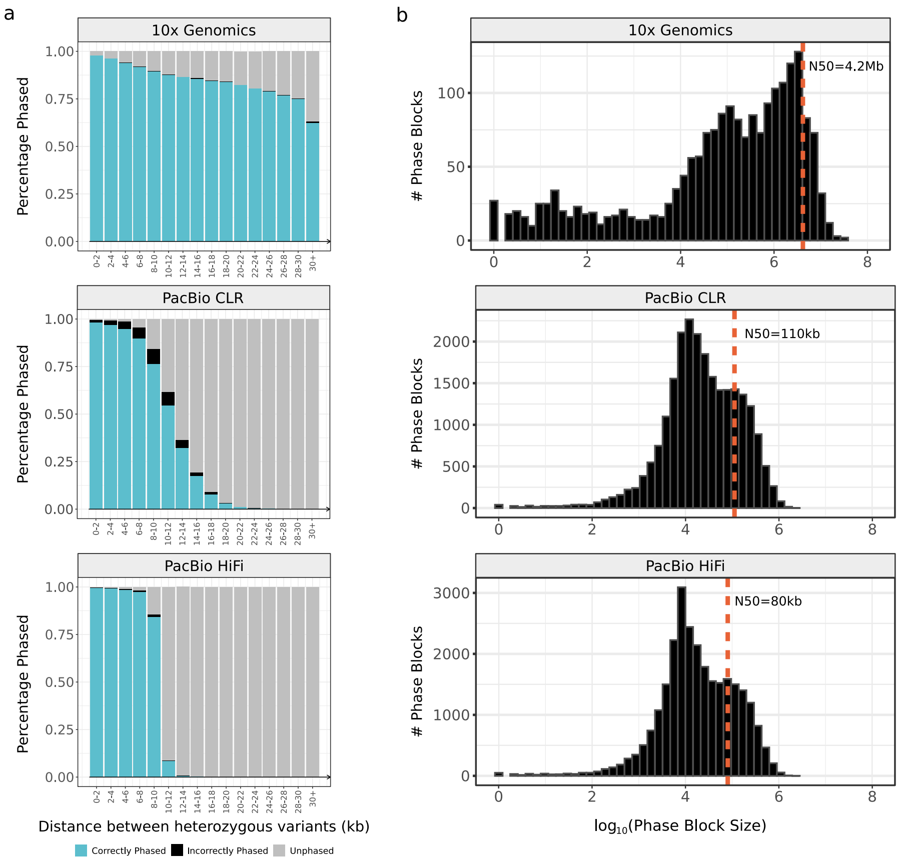


**Supplementary Fig. 7.** Phasing HG002 variants using different sequencing technologies. Variant phasing was performed using `Whatshap` for 10XG, PacBio CLR, and PacBio HiFi reads from HG002. Phased variants were compared against the GIAB high-confidence small variant truth set (v4.2.1). **a**, Breakdown of phasing accuracy based on the distance between pairs of neighboring heterozygous variants, binned by distance. Blue indicates phasing concordant with the truth set, black indicates discordant phasing, and grey represents unphased pairs. 10XG linked reads showed the best phasing performance across long distances (>30 kb) with high accuracy. PacBio CLR and HiFi reads showed phasing proportional to their average read length but with a longer tail and higher error rate for CLR. **b**, Distribution of phase block lengths ($\log_{10}$ scale) for each technology. N50 phase block lengths are shown as vertical dotted lines: 4.2 Mb for 10XG, 110 kb for PacBio CLR, and 80 kb for PacBio HiFi.

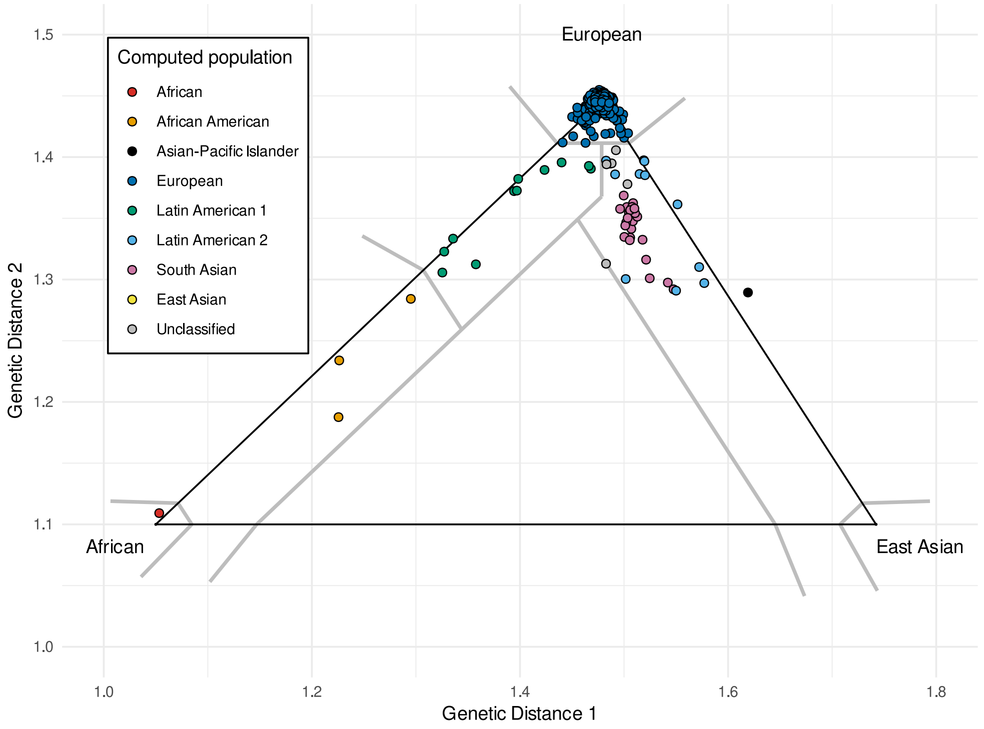


**Supplementary Fig. 8.** Genetic ancestry estimation for the full CF cohort sequenced with 10XG. Ancestry estimations for 564 individuals with CF were generated using `GRAF-pop` [22] based on 100,180 genome-wide SNPs called from 10XG reads. The cohort includes 101 individuals overlapping with the hybrid graph assembly set. Assigned ancestries were: European ($n$=506), African American ($n$=3), African ($n$=1), Latin American 1 ($n$=11), Latin American 2 ($n$=11), South Asian ($n$=25), Asian–Pacific Islander ($n$=1), and unclassified ($n$=6).

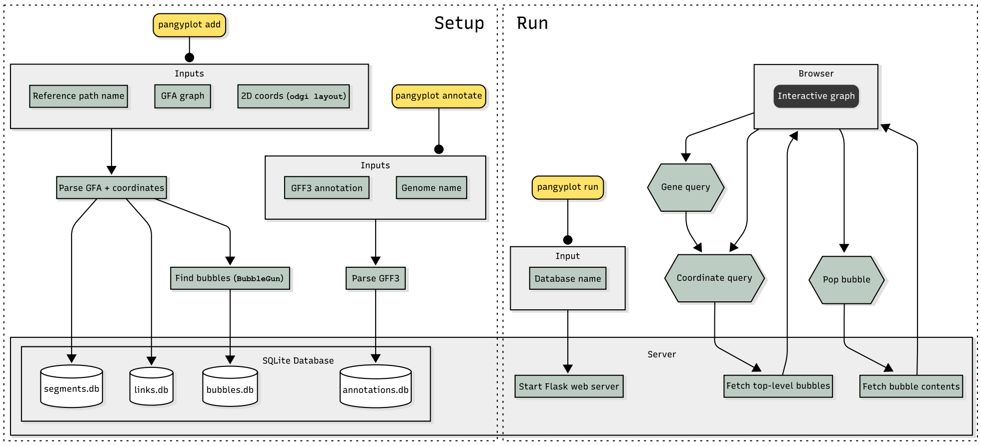


**Supplementary Fig. 9.** Flowchart of `PangyPlot` command-line subcommands for setting up and running the application. During setup, `pangyplot add` ingests a GFA file, a primary reference path name, and two-dimensional coordinates generated by `odgi layout`. `BubbleGun` functionality is integrated into `PangyPlot` for identification of bubbles and bubble chains. GFA data are indexed into multiple SQLite databases. Gene annotations can be indexed using `pangyplot annotate`, which loads a GFF3 file into a separate database. After setup, a Flask server can be launched with `pangyplot run` to respond to queries from the browser interface.

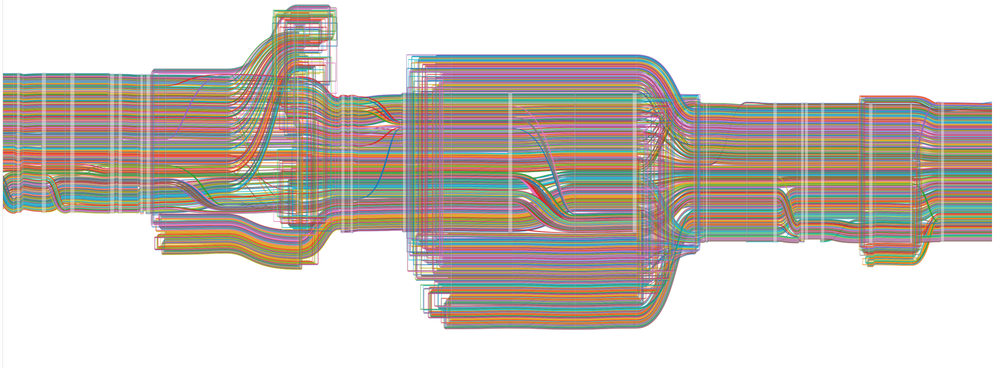

**Supplementary Fig. 10.** Sequence Tube Map visualization of a complex *CFTR* region. `Sequence Tube Map` [23] displays haplotypes as colored lines traversing sequence segments represented by white boxes. The figure shows a region within *CFTR* exhibiting reduced interpretability due to looping paths and visually entangled haplotype structures, highlighting the limitations of `Sequence Tube Map` in representing regions with complex variation.

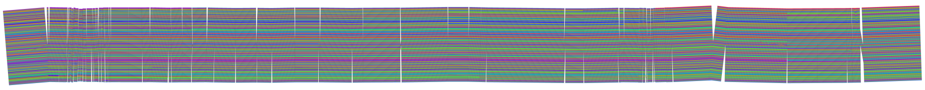

**Supplementary Fig. 11.** Two-dimensional `odgi layout` visualization of *CFTR*exons 10 and 11.

Two-dimensional layouts of variation graphs were generated using `odgi layout` followed by `odgi draw`. The figure shows a region encompassing *CFTR* exons 10 and 11. The high path density results in a tightly packed visualization where small variants are obscured.

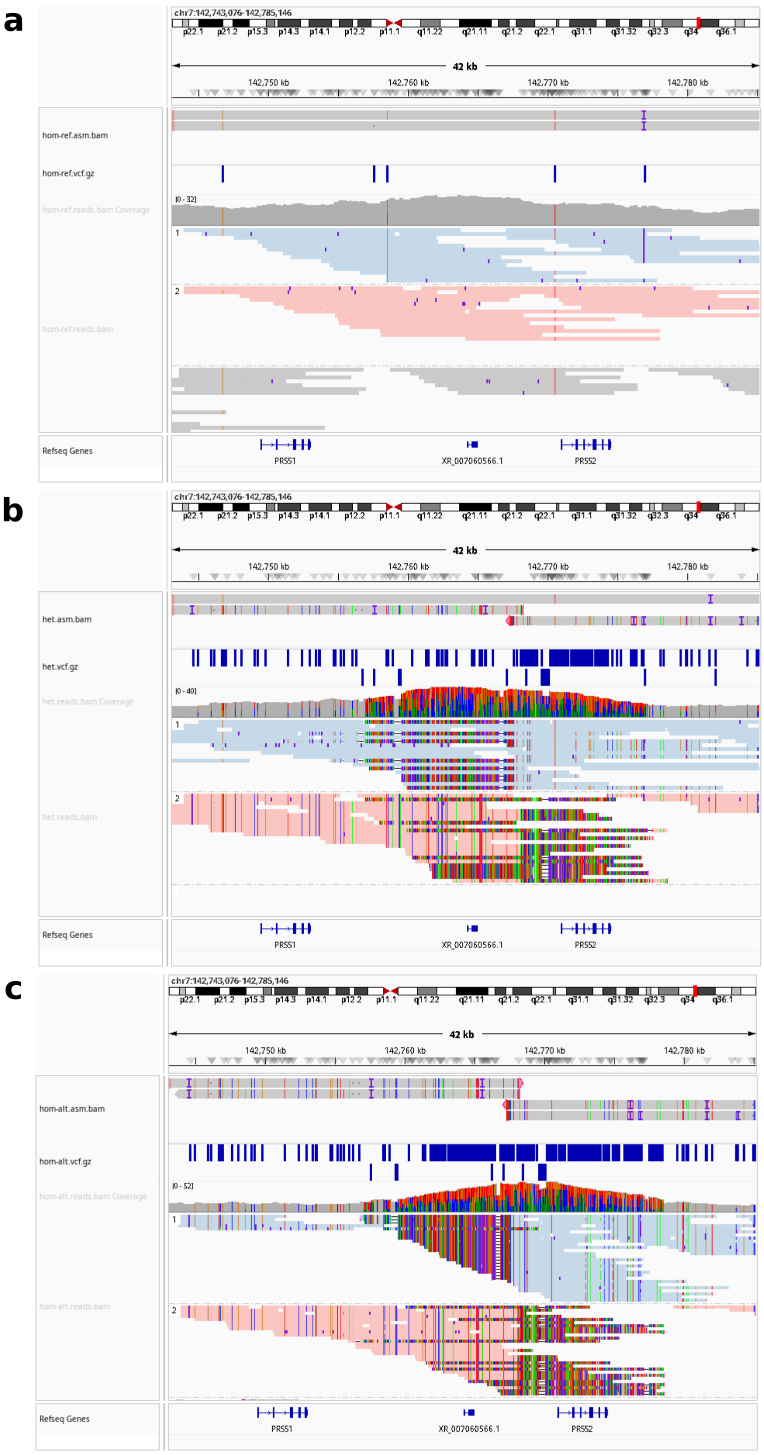


**Supplementary Fig. 12.** HiFi reads align poorly at the *PRSS1–PRSS2* locus due to reference bias. Alignments of PacBio HiFi data to the *PRSS1–PRSS2* locus on GRCh38 (3-copy allele) from three different individuals. Each panel includes (from top to bottom): a `Hifiasm` assembly aligned to GRCh38, a track showing variant calls from HiFi read alignment, a coverage track, and phased HiFi reads colored by haplotype (pale red and blue indicate phased reads; grey indicates unphased). Soft clipping appears as dense colored segments. **a**, 3-copy / 3-copy individual. Alignments are clean, phased correctly, and exhibit minimal reference bias; variant calling and coverage are consistent with the expected structure. **b**, 5-copy / 5-copy individual. Soft clipping is present with local coverage spikes, dense clusters of spurious variant calls, and phasing inconsistencies. **c**, 5-copy / 5-copy individual. All reads spanning the insertion region show soft clipping with highly inflated coverage and phasing errors. These alignment artifacts reflect strong reference bias when mapping to GRCh38. All three individuals had accurately assembled haplotypes in their `Hifiasm` assemblies.

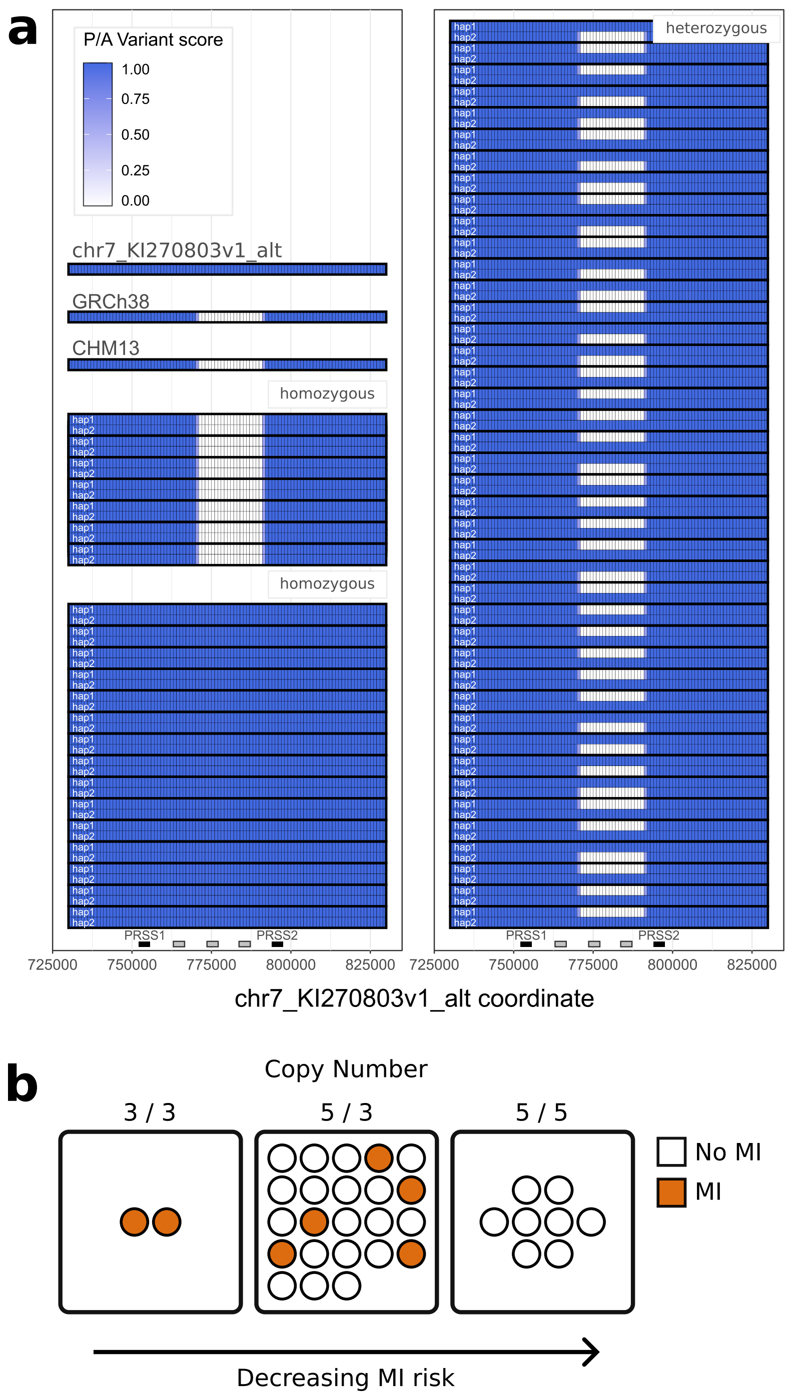


**Supplementary Fig. 13.** *PRSS1–PRSS2* CNV calls from HiFi assemblies are consistent with prior findings. **a**, Presence–absence variant plot from a 64-sample HiFi-based variation graph. Each cell represents a 1 kb window on the GRCh38 alternative contig KI270803.1; blue cells indicate presence of sequence, and white cells indicate absence. Pairs of rows correspond to an individual's haplotypes, with samples grouped by copy number genotype: 3-copy homozygous ($n$=7), 5-copy homozygous ($n$=15), and heterozygous ($n$=42). Gene annotations (black rectangles) and pseudogene annotations (grey rectangles) are shown. Observed allele frequencies of 44% (3-copy) and 56% (5-copy) match previously reported frequencies [24]. **b**, Lower copy number has been previously linked to increased risk of CF-related meconium ileus (MI) [24]. Here, individuals with known MI status (orange = MI, white = unaffected) show directionally consistent results, though the sample size is too small to demonstrate statistical significance.

**Supplementary Table 3.** Comparison of implementation details, strengths, and weaknesses of Sequence Tube Map, `odgi` (`viz` and `draw` subcommands), Bandage, and PangyPlot. *Chromosome-scale visualization in PangyPlot is a priority for future versions.*

| **Feature** | Sequence Tube Map | `odgi viz` (1D) | `odgi draw` (2D) | Bandage | PangyPlot |
|---|---|---|---|---|---|
| Required Format | vg | odgi | odgi | GFA | GFA |
| Style | Path-based, intersecting lines | Path-based, linear layout | 2D static image | Node-and-edge | Node-and-edge |
| Interactive | Yes | No | No | Yes | Yes |
| File Size Limitations | 5 MB | No specific limit | No specific limit | Must fit in memory | No specific limit |
| Layout | Real-time calculation | Based on graph sorting | Precomputed via `odgi layout` | Real-time calculation | Precomputed via `odgi layout` |
| Subsetting Required | Yes | No | Optional | Yes (for large graphs) | No |
| Implementation | Browser-based | Command line | Command line | Desktop application | Browser-based |
| Annotations | None | Possible to inject into graph | Possible via BED file | Requires external BED file | Built-in gene annotations |
| Query Options | Path coordinate-based | Range can be specified | – | Path coordinate, node ID, BLAST | Linear genome coordinates, gene-based |
| Setup | Install and deploy web app | Requires graph sorting | Requires graph sorting and layout computation | Minimal (load GFA in desktop app) | Layout computation, SQLite setup, app deployment |
| Small Variant | Yes | Not ideal | No | Yes | Yes |
| Structural Variant | Not ideal | Yes | Yes | Yes | Yes |
| Chromosome-scale | No | Yes | Yes | No | No* |
| Strengths | Visualizing haplotype distribution | Visualizing haplotype distribution, structural variants | Visualizing large-scale graph structure | Rich features, low setup overhead | Pangenome scalable, built-in annotations, physics engine |
| Challenges | Subsetting, locating regions of interest, complex variants | Highly dependent on sorting method | Tuning of drawing parameters to match graph scale | Memory limitation, manual layout optimization | Initial data preprocessing and indexing |

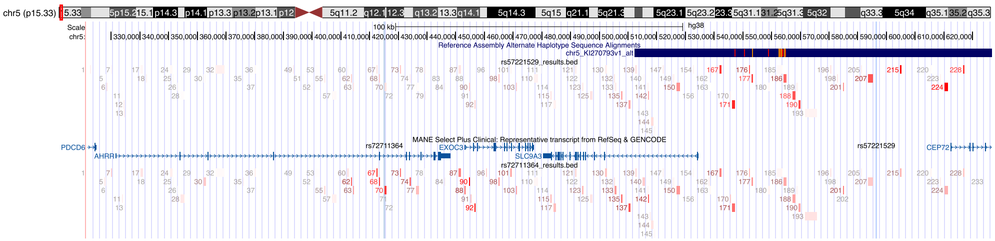

**Supplementary Fig. 14.** Genome browser view of repeat–SNP associations.

Genome browser visualization of the GRCh38 chromosome 5 subtelomeric region (chr5:311,504–626,711), highlighting tandem repeats from the Adotto catalog [25]. Tracks display repeats whose lengths are significantly associated with rs57221529 (upper) and rs72711364 (lower). Color intensity (red) reflects the significance of the logistic regression test ($p$-value), normalized to the most significant association per SNP. The middle track shows the positions of the two GWAS SNPs and nearby gene annotations.
